# Massively Parallel Polyribosome Profiling Reveals Translation Defects of Human Disease-Relevant UTR Mutations

**DOI:** 10.1101/2024.04.11.589132

**Authors:** Jia-Ying Su, Wei-Ping Li, Mijin Kim, Yu-Chi Chang, Hung-Lun Chiang, Yun-Lin Wang, Ang-Chu Huang, Yu-Tung Hsieh, Yi-Hsuan Chiang, Yen-Ling Ko, Bing-Jen Chiang, Cheng-Han Yang, Yen-Tsung Huang, Chien-Ling Lin

## Abstract

The untranslated regions (UTRs) of mRNAs harbor regulatory elements influencing translation efficiency. Although 3.7% of disease-relevant human mutations occur in UTRs, their exact role in pathogenesis remains unclear. Through metagene analysis, we mapped pathogenic UTR mutations to regions near coding sequences, with a focus on the upstream open reading frame (uORF) initiation site. Subsequently, we utilized massively parallel poly(ribo)some profiling to compare the ribosome associations of 6,555 pairs of wildtype and mutant UTR fragments. We identified 46 UTR variants that altered polysome profiles, with enrichment in pathogenic mutations. Both univariate analysis and the elastic net regression model highlighted the significance of motifs of short repeated sequences, including SRSF2 binding sites, as mutation hotspots that lead to aberrant translation. Furthermore, these polysome-shifting mutations exhibited considerable impact on RNA secondary structures, particularly for upstream AUG-containing 5’ UTRs. Integrating these features, our model achieved high accuracy (AUROC > 0.8) in predicting polysome-shifting mutations in the test dataset. Additionally, several lines of evidence indicate that changes in uORF usage underlie the translation deficiency arising from these mutations. Illustrating this, we demonstrate that a pathogenic mutation in the IRF6 5’ UTR suppresses translation of the primary open reading frame by creating a uORF. Remarkably, site-directed ADAR editing of the mutant mRNA rescued this translation deficiency. Overall, our study provides insights into the molecular mechanisms of UTR mutations and their links to clinical impacts through translation defects.

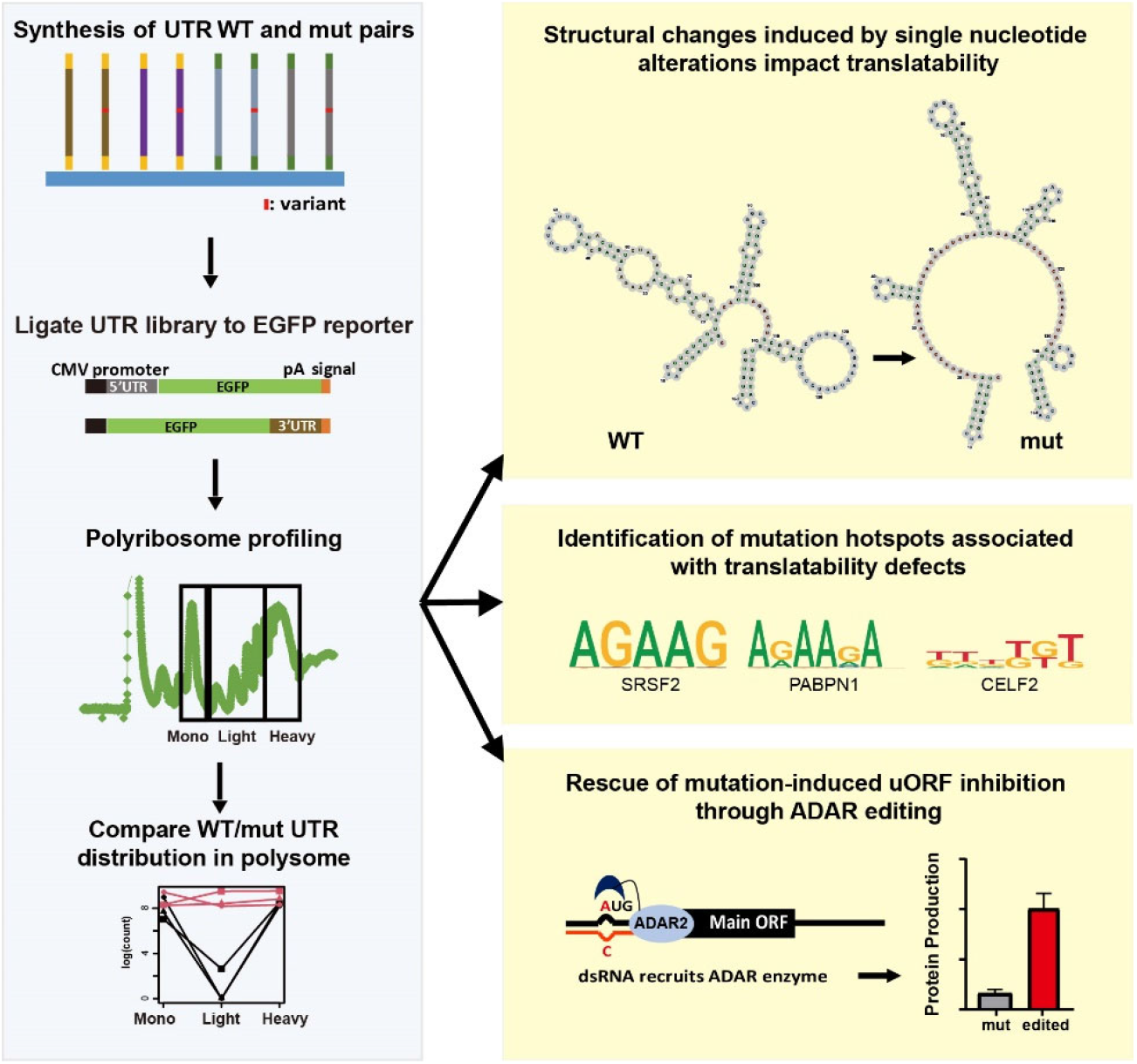

## INTRODUCTION

### Untranslated regions (UTRs) control translation efficiency

UTRs are crucial to gene expression regulation. Whilst mRNA coding regions determine protein sequence, the UTRs modulate protein abundance via, for instance, tuning stability and translation of mRNAs thereof. Many studies have shown that structural elements in 5’ UTRs are involved in translation regulation. An iron regulatory protein (IRP) that maintains iron homeostasis binds to an iron-responsive element (IRE) in the 5’ UTR of ferritin mRNAs, blocks recruitment of translation initiation factors, and elicits translation inhibition (Klausner et al., 1993; Muckenthaler et al., 1998). Internal ribosome entry sites (IRESs) are another *cis*-acting element of 5’ UTRs that control translation. IRESs in 5’ UTRs facilitate translation initiation by presenting ribosome-binding sites in a 5’ cap-independent manner (Komar & Hatzoglou, 2005). On the contrary, guanine-quadruplex motifs, which can adopt a non-canonical secondary structure in the 5’ UTR, are known to repress translation (Kumari et al., 2007; Serikawa et al., 2018). On the other hand, although 3’ UTRs lie far from the translation initiation site, they can regulate translation efficiency by controlling polyadenylation and binding of *trans*-acting factors. For instance, the poly(A) tail of mRNA can loop with its 5’ end, facilitating translation and ribosome recycling (Choe et al., 2018; Wells et al., 1998). Moreover, proteins and miRNAs that are deposited on 3’ UTRs can regulate various phases of translation including initiation, elongation, and termination (Mazumder et al., 2003).

### Massively parallel translation assays

To decode the regulatory potential of UTRs, we developed a massively parallel reporter assay (MPRA), in which UTRs of fixed length adjoin the coding sequence of a reporter. Similar approaches have been adopted to reveal the sequence-specific regulation and modular functions of numerous processes, including transcription, RNA splicing, translation, RNA localization, RNA decay and protein degradation in various species (Abell et al., 2022; Chiang et al., 2022; Griesemer et al., 2021; Jia et al., 2020; Litterman et al., 2019; Mikl et al., 2022; Niederer et al., 2022; Oikonomou et al., 2014; Rabani et al., 2017; Sample et al., 2019a; Savinov et al., 2021; Slutskin et al., 2018; Zhao et al., 2014). In some studies of RNA translation, reporter protein output, for example, intensity of the green fluorescent protein (GFP), has been used as a readout of regulatory activity. However, protein output essentially is the product of the interaction between RNA half-lives and its translatability. Therefore, for a more unequivocal study design, poly(ribo)some profiling has been used to estimate RNA translatability through evaluating their ribosome association status. MPRA based on polyribosome profiling technique has been applied to examine the translational efficiency of UTRs of interest. For instance, it has been shown previously that variations in 5’ UTRs have a greater impact on ribosome load than those in coding regions or 3’ UTRs (Leppek et al., 2022). MPRAs on 5’ UTRs have demonstrated that an upstream start codon (uATG) or open reading frame (uORF) can suppress primary ORF translation (Jia et al., 2020; Sample et al., 2019a). Structural hindrances, such as RNA G-quadruplexes (RG4) and stem-loops, can also suppress translation (Fay et al., 2017; Jia et al., 2020; Leppek et al., 2022). Additionally, at the sequence level, G-rich and C-rich elements are repressors of translation, whereas A-rich elements enhance translation (Jia et al., 2020; Niederer et al., 2022). Most massively parallel polysome profiling studies exclusively examined 5’ UTRs, likely due to the general assumption that 5’ UTRs play the major role in ribosome recruitment. On the other hand, MPRAs assessing fluorescent protein expression showed that binding sites for miRNAs and RNA-binding proteins (RBPs), particularly AU-rich elements, played a critical role in regulating the protein output (Oikonomou et al., 2014; Stoecklin et al., 2001; Wissink et al., 2016). Pumilio and AU-rich element-binding proteins, such as Tristetraprolin (TTP) and the embryonic lethal abnormal vision (ELAV) family, have been shown to influence protein expression (Barreau et al., 2005; Mayya & Duchaine, 2019; Van Etten et al., 2012). Nevertheless, whether these RBP or miRNA binding sites dampened protein level via impeding ribosome association was not corroborated by those MPRA studies.

### Precision diagnosis of UTR mutations

The genetic variants in UTRs may affect mRNA stability, translation, and localization, which in turn impact the flow of gene expression and hence result in phenotypical change and even pathogenesis. Prior studies have evidenced that even single nucleotide changes in a UTR can affect mRNA translation and/or transcript half-life in disease contexts. For example, single nucleotide substitutions at the 36^th^ position of the 5’ UTR or the 1723^rd^ position of the 3’ UTR of transforming growth factor β3 (TGFβ3; MIM# 190230) are associated with typical arrhythmogenic right ventricular cardiomyopathy (Beffagna et al., 2005). Another such example is a point mutation in the 3’ UTR of glutamine-fructose-6-phosphate transaminase 1 (GFPT1), which results in congenital myasthenic syndrome. GFPT1 is a rate-limiting enzyme for hexosamine biosynthesis, and a mutation in its 3’ UTR has been known to cause 90% reduction in protein production, potentially due to gain of a miRNA binding site (Dusl et al., 2015). All things considered, sequence integrity of UTRs has profound effects on gene regulation.

To systematically evaluate the impact of disease-relevant UTR variants on translation, we employed massively parallel ribosome profiling on both 5’ and 3’ UTRs to examine the effects of mutations on ribosome association. Through this strategy, we identified 46 high-confidence variants that caused ribosome shifting, and, intriguingly, their functional consequences converged on changes of RNA secondary structure and RBD binding motifs. Moreover, we investigated the molecular mechanism of a severe 5’ UTR suppressive mutation and developed an ADAR-based editing strategy to rescue the expression defect.

## MATERIAL AND METHODS

### Metagene analysis

All UTR and uORF variants were assigned using dbSNP version 151 (Sherry et al., 1999), with disease variants and UTR sources as described above. Definitions of uORFs were downloaded from TIS-db (Wan & Qian, 2014), and the respective genome coordinates were lifted from the Genome Reference Consortium hg19 to hg38 before analysis.

### Variant collection

Disease-related variants were collected from ClinVar (Landrum et al., 2016) and the HGMD database (Stenson et al., 2017). Variants located in the coding region or labeled as benign variants were excluded. UTRs were defined using NCBI RefSeq and ENCODE V27. Then, we intersected selected variant positions with the UTR regions by using BEDTools (Quinlan & Hall, 2010) to collect UTR variants.

### Construction of a UTR library of DNA templates

A UTR library of DNA templates was assembled by overlapping extension polymerase chain reaction (PCR) using Herculase II Fusion Enzyme (Agilent Technologies). Initially, the oligonucleotide library sequences (CustomArray Inc., USA) were double-strandized and amplified by PCR. After PCR clean-up (Qiagen), the UTR library amplicons were appended sequentially with EGFP CDS and CMV promoter sequences (derived from EGFP-N1 vector) by means of overlapping PCR. The assembled full-length DNA templates (CMVP-5’UTR-EGFP or CMVP-EGFP-3’UTR) were subjected to PCR clean-up for subsequent transfection. Primers are listed in Supplemental Table S1.

### Polysome fractionation and preparation for next generation sequencing

Polysome fractionation was conducted in accordance with a protocol published previously (Gandin et al., 2014) with slight modification. In brief, HEK293T cells cultured in 15-cm Petri dishes were grown to 60-70% confluency and then transfected with 4 µg of the UTR library of DNA templates (2 µg for each UTR library) using lipofectamine 3000 reagent (Invitrogen). Fourteen hours after transfection, the cells were incubated in culture medium containing 100 μg/ml cycloheximide (CHX) (Sigma) for 5 minutes.

Subsequently, the cells were washed twice with and then detached in ice-cold 1x PBS containing 100 μg/ml CHX, followed by centrifugation at 200 x *g* for 5 minutes at 4 °C. The entire procedure was conducted at a low temperature with the addition of 100 μg/ml CHX to preserve the polysome status. After discarding the supernatant, the cell pellet was re-suspended with 425 µl of hypotonic buffer (5 mM Tris-HCl (pH 7.5), 2.5 mM MgCl_2_, 1.5 mM KCl and 1x EDTA-free protease inhibitor cocktail (Roche)). Next, 5 μl of 10 mg/ml CHX, 1 μl of 1 M DTT, and 100 units of RNase inhibitor (Invitrogen) were added into the cell suspension, followed by vortexing for 5 seconds. Then, 25 μl of 10% Triton X-100 and 25 μl of 10% Sodium Deoxycholate were added into the suspension. After lysing the cells on ice for 5 minutes, the lysate was centrifuged at 16,000 x *g* for 7 minutes at 4 °C. The supernatant was collected and then layered onto a 5-50% sucrose gradient made of 5% and 50% sucrose solutions containing 20 mM HEPES (pH 7.6), 0.1 M KCl, 5 mM MgCl_2_, 10 µg/ml CHX, 0.1x EDTA-free protease inhibitor cocktail, and 10 units/ml RNase inhibitor, before undergoing ultracentrifugation with a SW41 Ti rotor (Beckman Coulter) at 36,000 rpm and 4 °C for 2 hours. Afterwards, the gradient samples were fractionated by fixed volume (500 µl/fraction), and all of the fractions were collected and subjected to RNA extraction (Direct-zol™ RNA MiniPrep Plus, Zymo Research). In addition to the indication of polysome profile derived from UV absorbance, the identity of each fraction was further verified through examining the distribution of 18S and 28S rRNAs among fractions via RNA electrophoresis.

Next, fractions corresponding to ribosome peaks were further categorized into three groups: monosome (Mono), low-density polysome (Light, 2-5 ribosomes), and high-density polysome (Heavy, ≥ 6 ribosomes). Then, the RNAs extracted from the fractions corresponding to each group were pooled together in an equal-fraction manner and subjected to reverse transcription using SuperScript IV First-Strand Synthesis System (Invitrogen). Thereafter, the UTR regions of mRNAs derived from our UTR library of DNA templates were amplified from the cDNA products using Phusion PCR Master Mix/GC Buffer (Thermo) and primers hosting Illumina sequencing adaptors and random tri-nucleotides for Illumina^®^ NextSeq paired-end 150 sequencing.

### High-resolution polysome profiling

For high-resolution translational profiling, gradient samples were classified into 10 distinct ribosomal states: free ribonucleoproteins (Free), 40S small subunits (40S), and individual polysome fractions (Pol1 to Pol8). Isolated RNA (500 ng) from each fraction was reverse-transcribed into cDNA using the SuperScript IV Reverse Transcriptase (Invitrogen) primed with custom oligonucleotides hosting a 3-nt fraction barcode and a 9-nt unique molecular identifier (UMI). Following reverse transcription, cDNA concentrations were measured to determine a baseline volume corresponding to 250 ng. To preserve the relative RNA abundance across gradient fractions, we adjusted the amount of cDNA used for PCR amplification based on the total RNA recovered from each fraction. Total RNA was calculated by multiplying RNA concentration by the pooled fraction volume (concentration × pooled volume), and the resulting values were normalized to the monosome (Pol1) fraction to determine the relative cDNA input for each fraction. Sequencing libraries were generated through a two-step PCR strategy using Phusion High-Fidelity DNA Polymerase (Thermo Fisher Scientific). First-round PCR was performed using target-specific primers to amplify the cDNA templates. The amplicons were purified using TOOLS Gel extraction and PCR cleanup Kit (Biotools) and subjected to second-round PCR using primers containing Illumina Nextera sequencing adapters (P5 and P7) (primer sequences are provided in Supplemental Table S2). Following secondary PCR, equal volumes of the amplicons across all 10 ribosomal fractions were pooled for the 5’ UTR and 3’ UTR libraries, respectively. The pooled amplicons were first concentrated and cleaned up using the same purification kit, followed by size selection on a 2% agarose gel. The target-sized library bands for the 5’ UTR and 3’ UTR pools were excised and purified using the QIAquick Gel Extraction Kit (Qiagen). The final dual-indexed 5’ and 3’ UTR libraries were quantified and subjected to paired-end 150 bp sequencing on the Illumina platform.

### RNA sequencing analysis

Raw sequencing data quality was evaluated using FastQC (v0.12.1). Nextera adapter sequences and low-quality bases were removed using Trimmomatic (v0.40) (Bolger et al., 2014) in paired-end mode with adapter clipping against the NexteraPE-PE adapter set (seed mismatch tolerance 2, palindrome clip threshold 30, simple clip threshold 10), a 4-base sliding window requiring a mean Phred score of at least 20, and a minimum read length of 70 bp; only read pairs in which both mates survived trimming were retained, and trimmed reads were reassessed with FastQC. Fraction-specific 3-nt barcodes and 9-nt UMIs (for high-resolution profiling only) were extracted from R2 reads and appended to the read names using umi_tools extract (v1.1.5) (Smith et al., 2017) with library-specific regular expressions. In both libraries the barcode and UMI cassette in R2 was anchored by fixed flanking sequences derived from the vector backbone or the EGFP reporter insert, and R1 carried a constant flanking sequence at its 5’ end; these constant sequences were removed before alignment, and read pairs that did not match the expected amplicon structure were discarded. Processed read pairs were aligned to the corresponding 5’ or 3’ UTR reference, consisting of the designed variant sequences with their flanking constant regions, using HISAT2 (v2.2.1)(Kim et al., 2019) with spliced alignment disabled and a stringent scoring scheme (using --mp 1,1 --rdg 1,1 and --score-min L,0,-0.7 --mp 50,25 -- rdg 15,12 --rfg 15,12 for low and high-resolution profiling respectively) that strongly penalizes mismatches and effectively prohibits gapped alignments, so that each read is assigned only to its exact variant of origin; a single best alignment was reported per read (-k 1). Alignments were coordinate-sorted and indexed using SAMtools (v1.19) (Danecek et al., 2021), and only uniquely mapped reads (MAPQ = 60) were retained. PCR duplicates were collapsed using umi_tools dedup in paired-end mode with the unique method, which merges read pairs sharing an identical UMI and alignment coordinates without UMI error correction. Deduplicated alignments were demultiplexed into the 10 ribosomal fractions (Free, 40S, and Pol1 to Pol8) according to the fraction barcode in each read name; reads carrying unrecognized barcodes were discarded. Finally, per-variant read counts for each fraction were obtained by counting aligned reads overlapping the full length of each reference sequence using BEDTools multicov (v2.31.0) (Quinlan & Hall, 2010), yielding a variant-by-fraction count matrix for downstream translational profiling.

### Statistical method for detecting the effect of mutations on polysome profiles

To determine variants that changed polysome distribution, we used read counts in polysome fractions to build a negative binomial generalized linear model with a log link function for each pair of oligonucleotides as follows:

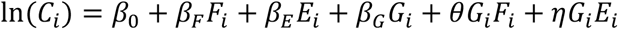

where *C*_i_ are read count values of the ith replicate, *F*_i_ represents the three polysome fractions (Mono, Light and Heavy), *E*_i_ represents biological replicates, and *G*_i_ represents WT or mutant.

We tested the significance of *θ* to determine the effect of the variant. To avoid potential batch effects, we further excluded candidates with significant 5 or *β*_E_ (*p* < 0.05).

All statistical analyses were performed using R language (version 4.2.2). Linear models were constructed using the *glm.nb* function in R.

### Construction of DNA templates for luciferase assay

Linear constructs (CMVP-5’UTR-FFLuc or CMVP-FFLuc-3’UTR) for luciferase assay were assembled by overlapping extension PCR using Phusion DNA Polymerase (Thermo). Initially, DNA oligos (Integrated DNA Technologies Inc.) of selected UTR candidate sequences were double-strandized and amplified by PCR. Afterwards, the purified UTR amplicons were appended with firefly luciferase CDS (derived from pGL3-Basic vector) and the CMV promoter sequence (derived from EGFP-N1 vector) sequentially via overlapping PCR.

### Luciferase assay, RNA extraction, and RT-qPCR

Fourteen to fifteen hours after co-transfection with 500 ng of linear constructs (CMVP-5’UTR-FFLuc or CMVP-FFLuc-3’UTR) and 100 ng of pRL-TK vector, HEK293T cells cultured in a 12-well plate were harvested and processed according to the instruction of Dual-Glo Luciferase Assay System (Promega) with minor modification. In brief, after removing media and washing once with PBS, the transfected cells were detached and then lysed in 180 µl of Dual-Glo reagent, before being subjected to moderate shaking on an orbital shaker at room temperature for 10 minutes. The lysate was then transferred to a new tube and underwent low-speed centrifugation to remove the cell debris. Then, 50 µl of clear supernatant was transferred to 96-well black plates to measure firefly luminescence with EnSpire Multimode Plate Reader (PerkinElmer) or SpectraMax Paradigm reader (Molecular Devices) and the software SoftMax Pro. Subsequently, 50 µl of Dual-Glo Stop & Glo Reagent was added into each well. After incubation at room temperature for 10 minutes, *Renilla* luminescence was measured as above.

Relative luminescence units (RLUs) were calculated by dividing the firefly luciferase readouts by those of the transfection control, i.e., *Renilla* luciferase. To evaluate translation efficiency, the RLUs were normalized to the transcript level measured by real-time PCR. Apart from those specifically annotated herein, all data is shown as the fold-change of the mutant relative to wildtype.

To calibrate the luciferase assay results against cognate mRNA expression level, total RNAs were extracted from the cells subjected to identical transfection conditions with TRIzol reagent (Invitrogen). Total RNA was extracted with Direct-zol RNA Miniprep kits (Zymo Research), before being subjected to reverse transcription. Then, cDNA was reverse-transcribed from 1 µg of RNA using 2.5 µM random hexamer with 1 µl (200U) SuperScript IV Reverse Transcriptase (Thermo Scientific). The reaction mixture was incubated at 24°C for 10 min to allow primer annealing, followed by incubation at 50°C for 10 min for reverse transcription. Subsequently, the reaction was terminated by incubating at 80°C for 10 min to inactivate the reverse transcriptase. Finally, the cDNA products were subjected to real-time PCR with Fast SYBR Green Master Mix (QuantStudio 12K Flex system, Applied Biosystems) to determine the expression level of firefly luciferase relative to that of *Renilla* luciferase.

### Motif enrichment analysis

The motif analysis include RBP position weight matrix (PWM), K-mer and miRNA seed sequence. RBP PWM definitions were downloaded from the ATtRACT database (Giudice et al., 2016). For each UTR sequence, all possible motif PWMs were determined using the R package Biostrings (Pagès H, 2022) and the *matchPWM*() function with the parameter min.score = 95%. To summarize the contribution of each RBP, PWMs predicted for the same RBP gene were summed. K-mer composition was determined in Biostrings using the *oligonucleotideFrequency*() function, with K-mer lengths of 1 to 7. miRNA binding sites were predicted in TargetScan 7.0 (Agarwal et al., 2015), with high-confidence predictions (8mer, 7mer-m8, 7mer-1A) for each miRNA being summed to represent counts of the seed sequence.

To find motifs enriched in UTRs of interest, motif distribution in relation to UTRs was analyzed by Fisher’s exact test using the *fisher.tes*t() function in R (R-Core-Team, 2021) according to the matrix:

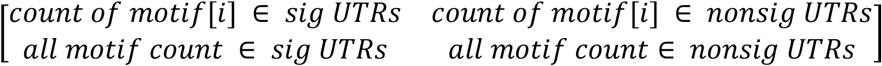

To examine motif changes in relation to polysome shifting, motifs disrupted by variants and the resulting shift in polysomes were analyzed by using the *fisher.tes*t() function in R according to the matrix:

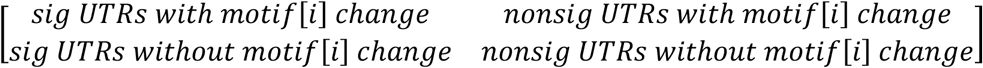

### shRNA-mediated knockdown

For shRNA expression, pLKO.1 vectors containing an shRNA sequence (5’-AGTTGTGTAGCAGTTGAGTAA-3’) targeting the human *SRSF2* 3’ UTR or a non-targeting scrambled control (5’-CCTAAGGTTAAGTCGCCCTCG-3’) were obtained from the RNAi Core, Academia Sinica, Taiwan, and lentiviral particles were produced using HEK293T cells by following standard procedures.

The HEK293T cells were initially seeded at a density of 5×10^5^ cells per well in 6-well plates. Four hours after seeding, shRNA-expressing lentivirus was introduced into the HEK293T cells at a multiplicity of infection (MOI) of 5. Then, the virus-containing medium was replaced after 24-hour incubation with 2 ml of DMEM supplemented with 10% FBS. Following an additional 24-hour incubation, the culture medium was supplemented with 2.5 μg/ml puromycin for selection purposes. The cells underwent a 4-day puromycin selection with one round of medium refreshment. The cells were then incubated in medium with 5 μg/ml puromycin for an additional day. After the selection process, the cells were transferred to a 10 cm dish for expansion and subsequent experiments.

### Plasmid construction

For the firefly luciferase reporter plasmids, full-length UTRs were PCR-amplified from HEK293T complementary DNA (cDNA) with Phusion High-Fidelity DNA Polymerase (Thermo Scientific). Restriction enzyme cutting sites (*HindIII* and *NcoI* for 5 ‘UTRs, *XbaI* for 3’ UTRs) were attached for digestion with Thermo Scientific FastDigest kit and for cloning with Rapid DNA Ligation kit (Thermo Scientific) into the pGL3-Promotor (Promega) backbone. Mutants were generated by mutagenesis PCR from wildtype constructs and the same cutting sites as for cloning were adopted.

For motif validation, the 5’ UTR sequence of *MUTYH* (mutY DNA glycosylase:c.-43∼-157) and the 3’ UTR sequence of *MEFV* (Mediterranean fever:c.*17∼131) were used, respectively, as the backbone of wildtype UTRs. Motifs were inserted into the *Xba1* site of the *MUTYH* 5’ UTR and the *Nhe1* site of the *MEFV* 3’ UTR.

For IRF6 ar151-expressing plasmids, the 151 bp antisense sequence with a mismatch in the center was PCR-amplified from wildtype luciferase plasmids. The PCR product was inserted into the pHE vector (BIOTOOLS) and transcription was driven by the human U6 promoter.

### UTR secondary structure prediction

UTR secondary structures were predicted and plotted using the forna website (http://rna.tbi.univie.ac.at/forna/) (Kerpedjiev et al., 2015). Free energy was estimated using the viennaRNA package (Lorenz et al., 2011) and the *RNAfold* command with the default setting.

### Elastic net model of polysome-shifting UTR variants

To develop a prediction model for determining whether a genetic variant affects mRNA translation, we employed a logistic regression model with elastic net regularization for feature selection. The curated features included free energy, structure rate, GC content, k-mer, G-quadruplex (RG4), conservation level, miRNA binding motif, PUM motif, RBP motif, and ARE. Before feature selection, the features were narrowed down using multiple logistic regression models to regress the significance of variants (Yes/No) on each feature, with a *p-value* < 0.01 serving as the criterion for identifying potential candidates. Subsequently, the dataset was divided into two groups, comprising training and testing sets in a 2:1 ratio, respectively. The model is represented as follows:

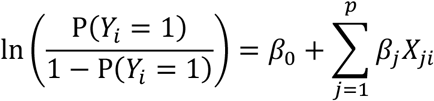

where *Y*_i_ = 1 is the ith mt sequence that significantly affects mRNA translation, and *X*_ji_ is the feature *j* in the *i*th sequence. The predictive features were standardized using Z-score transformation to achieve unit standard deviation, thereby ensuring comparability of coefficients. Elastic net regularization minimizes 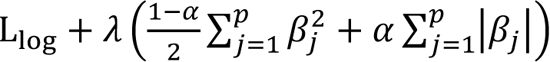, where the negative log likelihood is:

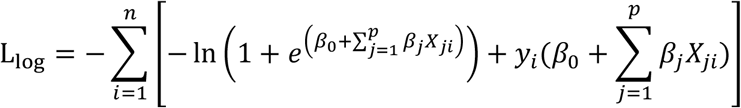

We applied 10-fold cross-validation on the training set to optimize the tuning parameters (λ and α) for the elastic net model. Following this, predictions were made on the testing set to evaluate the model’s performance, and the area under the receiver operating characteristic (AUROC) was calculated. Feature selection and AUROC calculations were conducted using the R packages ‘caret’ (v.6.0.94) and ‘precrec’ (v.0.14.4) (Kuhn, 2008; Saito & Rehmsmeier, 2017), respectively, with all analyses performed on R version 4.3.3. The 95% confidence intervals for the AUROC values were estimated through bootstrap resampling with 1,000 iterations.

### Validation of the elastic net model

To further validate the predictive performance of our elastic net model, we conducted external validation using an independent dataset published by Sample et al. (2019). This dataset comprises curated human 5’ UTR sequences that were systematically evaluated using a massively parallel translation assay to assess their effects on translation efficiency (Sample et al., 2019a). The dataset was obtained from the authors’ publicly available repository (https://github.com/pjsample/human_5utr_modeling).

Prior to analysis, rigorous quality control measures were implemented to ensure data reliability. Sequences were included only if they met the following criteria: an average read count of at least 50 reads per polysome fraction and a minimum of 10 reads in each individual fraction. Since our predictive model outputs binary classifications (significant effect versus no significant effect), we established a threshold to categorize the continuous mean ribosome load (MRL) change values from the Sample et al. dataset. Variants with an absolute log₂ mean ribosome loading (MRL) change of ≥1.0 (corresponding to the “log_obs_diff” column in their dataset) were classified as having significant translational impact, while those below this threshold were considered non-significant. Model performance on the external dataset was evaluated by calculating the AUROC with 95% confidence interval using the same methodology described previously.

It should be noted that external validation for the 3’ UTR predictive model could not be performed due to the absence of suitable independent datasets with comparable experimental design and quality standards.

### Translation efficiency

Translation efficiency (TE) was modelled for wild-type 5’ UTR sequences in HEK293T cells and measured in three biological replicates. For each sequence, reads were summed over the polysome fractions F2–F8 and over the ribosome-free and 40S fractions. A replicate was used for a sequence only if that sequence had at least one read in each of the two pools. Sequences with fewer than two usable replicates were excluded. TE was the mean of the per-replicate log₂ ratios,

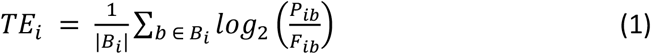

where *P*_ib_ and *F*_ib_ are the polysome and the ribosome-free plus 40S counts of sequence *i* in replicate *b*, and *B*_i_ is the set of usable replicates for that sequence. Sequences whose TE deviated from the mean by more than three standard deviations were excluded as outliers.

A ratio estimated from more reads is more precisely determined. Each sequence was therefore weighted by the square root of its mean total count, rescaled to unit mean,

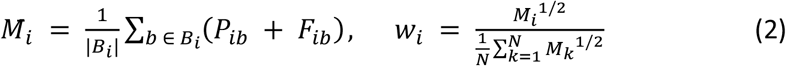

The weight depends on total depth alone and so carries no information about TE itself. The square root was used in place of the count so that the deepest sequences do not dominate the fit. The same weights were used at every step below.

### Feature selection

Many features describe much the same thing and are strongly correlated with one another, which makes feature selection unstable. Features were therefore reduced to a set of representatives, as described previously (Su et al., 2025). All features were clustered using single-linkage hierarchical clustering with the distance metric defined as one minus the absolute value of the Spearman correlation coefficient. Cutting the tree at a given height divides the features into clusters. Within each cluster, the influence of every feature on translation efficiency was examined using a weighted simple linear regression model, and the feature with the greatest influence became the representative of that cluster. Raising the height merges clusters and so leaves fewer representatives. The height was raised until every representative had a variance inflation factor below 5. The clusters, the univariate effect of every member and the representative selected from each are provided in Supplemental Table S3.

### Model building and evaluation

We performed a 2:1 split of the data into training and testing sets to validate the robustness of the model. The penalised regression and the stability selection below were carried out on the training set alone. The testing set was used only to compare predicted with observed translation efficiency.

LASSO regression was performed on the representative features, with the sequence weights of Equation 2,

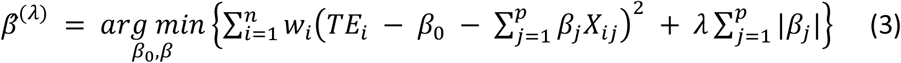

where *X*_ij_ is the standardised value of representative feature *j* for sequence *i*, *p* the number of representatives and *n* the number of sequences. λ was chosen by ten-fold cross-validation, as the value giving the smallest mean cross-validated error.

The set of features that a single LASSO fit selects can change appreciably when the data change. Features were therefore chosen by stability selection (Meinshausen & Bühlmann, 2010). One hundred subsamples of half the training sequences were drawn. In each subsample the weighted LASSO of Equation 3 was fitted with its own λ, and the features with non-zero coefficients were recorded. The selection probability of a feature is the fraction of subsamples in which it was selected,

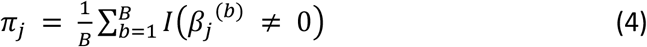

with *B* = 100 subsamples, β_j_^(b)^ the LASSO coefficient of feature *j* in subsample *b*, and I(·) the indicator function. Features with π_j_ > 0.7 were retained, a threshold in the range recommended by Meinshausen & Bühlmann (2010). The LASSO was used only to decide which features enter the model. Its coefficients are not reported.

The retained features were refitted on the training set by weighted linear regression,

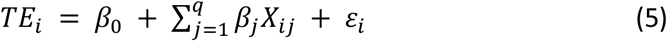

where *q* is the number of retained features and the regression was weighted by *w*_i_ as in Equation 2. The coefficients of this model are reported rather than the penalised ones, with their 95% confidence intervals. The predicted translation efficiency of any sequence, wild type or mutant, is

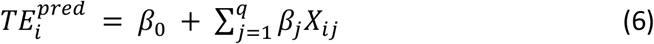

Predicted and observed translation efficiency were compared on the testing set by Spearman and Pearson correlation. Every prediction described below was made with this model.

### Validation against luciferase reporters

Three comparisons were made. First, translation efficiency was predicted for eight wild-type 5’ UTR constructs whose translation had been measured by luciferase assay in two replicates, and compared with the log₂ of the replicate mean. Second, for each variant group measured in both genotypes, the effect of the variant in the MPRA was ΔTE = TE(mutant) − TE(wild type). This was compared with the luciferase log₂ ratio of mutant to wild type. Third, the same comparison was made with the change the model predicts for that variant. Each comparison was made for library constructs, for full-length constructs and for the two pooled, and was summarised by the Spearman correlation coefficient.

### Software and reproducibility

Analyses were carried out in R version 4.6.1. Penalised regression used glmnet (version 5.0), with the penalty of each subsample chosen by its cv.glmnet function. Variance inflation factors were computed with car (version 3.1-5). Figures were produced with ggplot2 (version 4.0.3) and patchwork (version 1.3.2), and sequence logos with ggseqlogo (version 0.2.2). Every random draw is governed by a fixed prime seed assigned per partition. The subsamples and the split are therefore reproducible. *P* values of correlations were obtained with cor.test.

### Western blotting

To validate translation activity on the mutant-generated upstream AUG, HEK293T cells were cultured in 6-well plates. After 24 hours, the cloned reporter plasmids and the control pEGFP-N1 were co-transfected by Lipofectamine 3000 (Invitrogen) at a ratio of 3:1 (2.5 μg in total). At 24 h post-transfection, cells were lysed in RIPA buffer (Sigma-Aldrich) supplemented with cOmplete Protease Inhibitor Cocktail (Roche). The solution was transferred onto ice for incubation for 5 minutes and then centrifuged at full speed for 10 minutes at 4 °C. Finally, the supernatant was transferred to a new tube. After determining protein concentration by Bradford assay (Bio-Rad), 30-80 μg of protein sample was loaded into each well. Protein samples separated by sodium dodecyl sulfate-polyacrylamide gel electrophoresis were transferred to PVDF membrane for antibody incubation and ECL detection. Immunoblotting of IRF6: primary antibody against IRF6 (Cell Signaling Technology, Cat# 6948, dilution: 1:1000) and secondary antibody (Jackson ImmunoResearch, Cat#: 111-035-003, 1:5000); immunoblotting of tubulin: primary antibody against tubulin (Sigma, Cat#: T6793, 1:2000) and secondary antibody (Cell Signaling Technology, Cat#: 7076, 1:5000); immunoblotting of firefly luciferase: primary antibody against luciferase (Abcam, Cat#: ab185924, 1:1000) and secondary antibody (Abcam, Cat#: ab6721, 1:5000); immunoblotting of GFP: primary antibody against GFP (ProteinTech, Cat#: HRP-66002, 1:10000).

### ADAR editing

HEK293T cells were cultured in 24-well plates. After 24 hours, 200 ng of the reporter plasmids, 20 ng of the pRL-TK plasmids, 150 ng of the ar151 plasmids, and 150 ng of the ADAR2 overexpression plasmids were co-transfected with Lipofectamine 3000 (Invitrogen). Luciferase assays and evaluations of mRNA levels were performed as described above after 40 hours of transfection.

To evaluate the editing efficiency of IRF6 ar151, the cDNA synthesized for qPCR analysis was PCR-amplified with primers targeting 151 bp of the 5’ UTR. PCR products were purified with QIAquick PCR Purification Kit (QIAGEN) according to the manufacturer’s protocol. Elution products were subjected to Sanger sequencing to analyze editing efficiency.

## RESULTS

### Meta-analysis of disease-relevant UTR variants

It has been estimated that ∼3.7% of disease-relevant genetic variants arise in UTRs (MacArthur et al., 2017; Steri et al., 2018). To investigate if there is a positional effect of disease-relevant UTR variants, we compared the overall distribution of disease-associated single nucleotide polymorphisms (SNPs) in ClinVar (Landrum et al., 2016) to over a million human UTR SNPs in the dbSNP archive, which revealed enrichment for disease-relevant variants near mRNA coding regions in both 5’ and 3’ UTRs (**Fig. 1A**), indicating that the near-coding region of UTRs is a critical regulatory region for gene expression. Although we did not detect enrichment for disease-relevant 5’ UTR variants near the transcription start site, they did exist at uORF start sites (**Fig. 1B**). This specific localization of disease-relevant UTR variants evidences a trend in the positioning of functional elements within UTRs.

**Figure 1.**
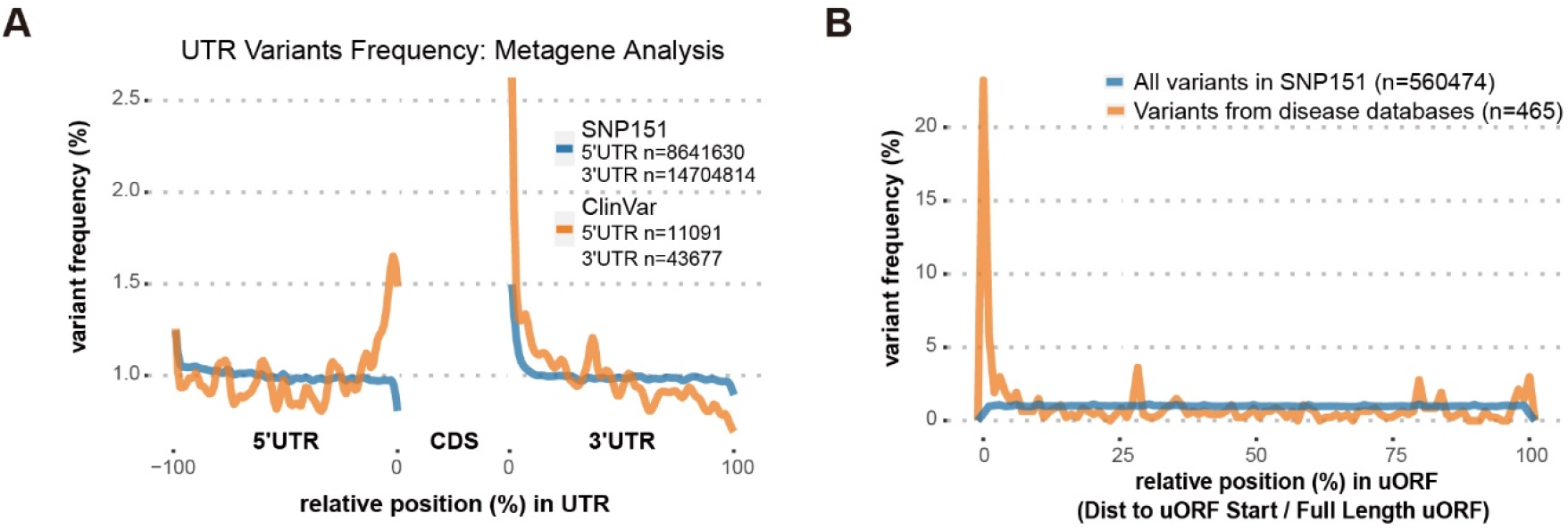
Enrichment of UTR variants relative to open reading frames. **(A)** Metagene analysis of native human UTRs (blue line) or disease-relevant variants reported in ClinVar (orange line). Note the enrichment of UTR variants near the coding region. **(B)** Metagene analysis of all native human variants (blue line) or disease-relevant variants reported in ClinVar and HGMD (orange line) in relation to uORFs. Note the distinct enrichment of disease-relevant variants at the uORF start site.

### Massively parallel poly(ribo)some assays on human disease-relevant UTR mutations

Although many disease-associated UTR variants are located in functional elements, their pathogenic mechanisms remain unknown. To examine if these variants directly affect protein output, we first sought to determine the impact of these variants on translation. To systematically examine the altered translation efficiency by UTR variants, we implemented massively parallel polysome assays to examine the effects of variants on polysome association (**Fig. 2A**). First, we extracted 6,555 disease-associated UTR variants, including SNPs and indels shorter than 15 nucleotides (nt), from the Human Gene Mutation Database (HGMD) and ClinVar, followed by multiplex oligonucleotide synthesis for selected UTRs. We prioritized UTR variants designated as pathogenic or likely pathogenic in the database. Consequently, all annotated pathogenic and likely pathogenic UTR variants reported by the time of data collection were included in the library. Additionally, due to the constraint of synthesis length (∼150 nt), we synthesized the segment of selected UTRs (∼115 nt) in which variants were located at the center to maximize the possibility of including critical regulatory elements. Moreover, UTR fragments of both alternative and reference alleles were synthesized in parallel for case-control comparison. In cases where multiple alternative alleles existed at the same position, they were paired with the same reference allele to prevent duplication in the library. The entire library of UTR oligonucleotides (UTR library) was subsequently ligated upstream or downstream of an enhanced GFP (EGFP) coding region, along with a CMV promoter and a common UTR sequence on the opposite end. Cells transfected with the UTR library were treated with cycloheximide 14 hours post transfection and then subjected to polysome fractionation (see Methods). By matching the UV absorbance profile of fractions to rRNA abundance revealed by gel electrophoresis of RNA samples (total RNA extracted from polysome fractions), we were able to estimate the number of bound ribosomes in the collected fractions. RNA samples from fractions containing 1 ribosome, 2-5 ribosomes, and 6 or more ribosomes were pooled, respectively, and are referred to hereafter as the Mono, Light and Heavy groups.

**Figure 2.**
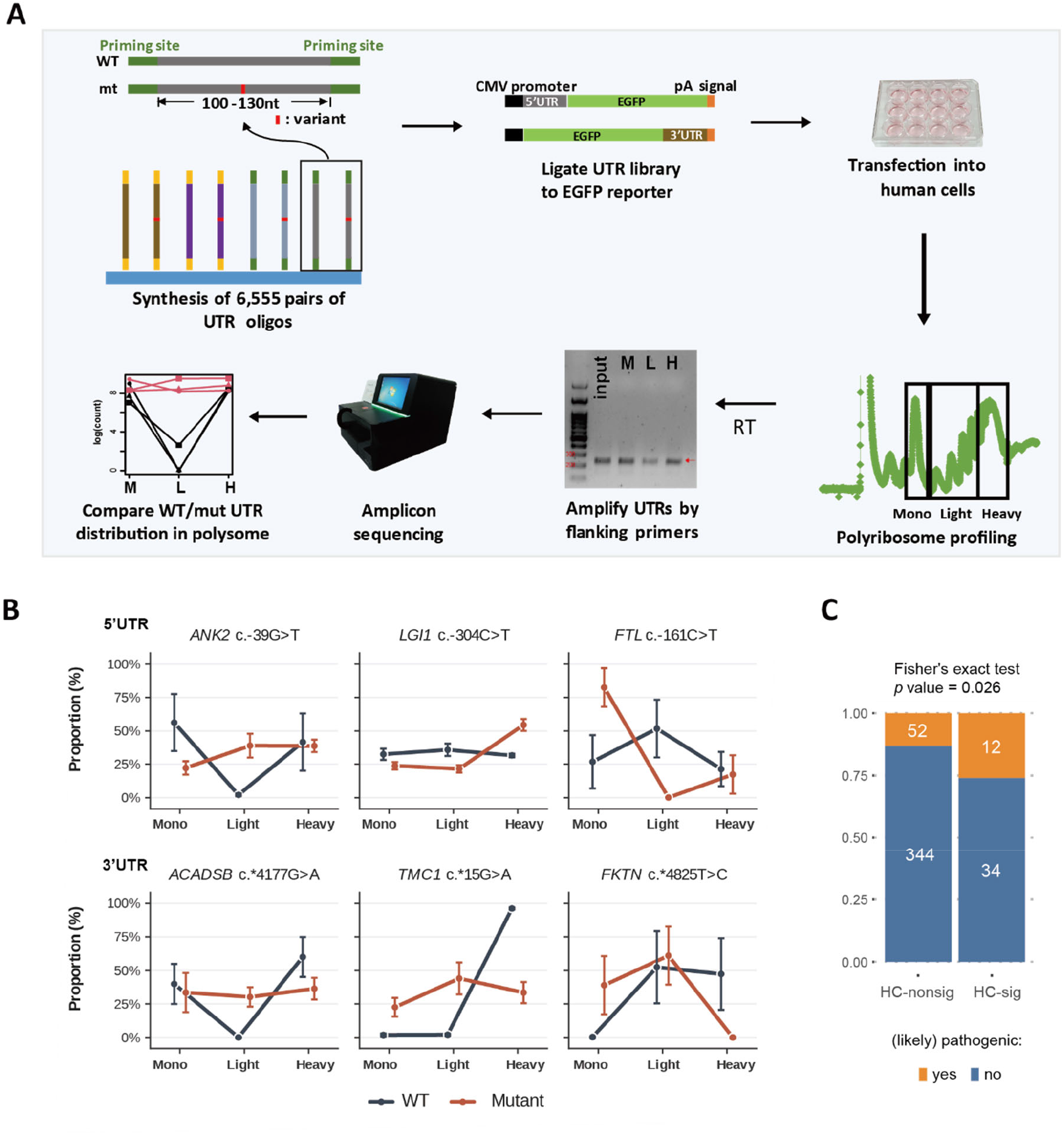
Massively parallel polysome assays. **(A)** UTR fragments were synthesized in bulk and ligated with EGFP reporters for expression in human cells. The reporter-expressing cell lysate was fractionated into monosome, light (2-5 ribosomes) and heavy (6+ ribosomes) polysome-containing fractions, and the UTR library was then amplified for amplicon sequencing. Mutations that altered relative distribution among fractions were defined as significant polysome-shifting mutations. **(B)** Examples of UTR pairs that significantly differed in polysome distribution. **(C)** Enrichment of pathogenic variants among HC-sig polysome-shifting UTR variants.

UTR library constituents were then reverse-transcribed and PCR-amplified from the three groups of RNA samples, before undergoing next-generation sequencing. To evaluate the effects of disease-associated variants on the distribution of corresponding transcripts across the Mono, Light and Heavy groups, we fit a negative binomial generalized linear model to read count for each UTR pair (wildtype/WT and mutant/mut) in three fraction groups while accounting for batch effects of the three experimental replicates (see Methods). The model primarily focused on discerning the distribution pattern of reads across three polysome fractions between UTR pairs, without considering differences in the expression levels of the UTR species. As a result, the analysis was unaffected by any perturbation of UTR sequence at the transcriptional level and did not account for changes in stoichiometry between UTRs and regulatory factors due to variations in expression levels. Consequently, we identified 483 high-confidence UTR pairs, comprising 396 pairs that did not alter translation (172 and 224 for 5’ UTRs and 3’ UTRs, respectively; collectively denoted by high-confidence nonsignificant or ‘HC-nonsig’ UTR variants), and 19 5’ UTR and 27 3’ UTR variants that significantly affected polysome distribution (**Fig. 2B, Supplemental Figs. S1, S2 and Table S4**; collectively denoted by high-confidence significant or ‘HC-sig’ UTR variants). Pearson correlation analysis revealed R coefficients ranging from 0.59 to 0.71 for the mut-to-WT transcript ratios across three independent experiments **(Supplemental Fig. 3)**.

Although a very small proportion of UTR variants are categorized as pathogenic, we observed that HC-sig variants are enriched with known pathogenic variants relative to the HC-nonsig variants (**Fig. 2C**), suggesting a contribution of translation-altering UTR variants to pathogenicity. Notably, we found that these translation-altering UTR variants are particularly prevalent in neurological and developmental disorders **(Supplemental Table S5)**.

Even with the high confidence dataset, we did not intend to immediately translate the altered polysome profile into an increase or decrease in translation efficiency, as the direction of the shift was not readily evident. Additionally, sedimentation in the sucrose gradient may have been partially affected by heavy particles other than ribosomes. Subsequently, we verified the effects of individual HC-sig variants on translation via luciferase reporter assay.

### Validating the translation defects of mutant UTRs by luciferase reporter assay

The variants significantly altering the polysome profile were then individually validated through high-sensitivity luciferase reporter assays (**Fig. 3A**). To this end, we resynthesized both the variant and corresponding wildtype alleles in the same library format - 115-nt native UTR segments centered on the variant and flanked by 20-nt priming sites. These UTRs were then cloned upstream (5’) or downstream (3’) of the firefly luciferase coding sequence, depending on their genomic location. As the initial library design, the test UTR segment differs from their WT counterparts by only one nucleotide, while a shared short UTR fragment is present on the opposite end of the coding sequence to ensure consistency across constructs (**Fig. 2A**). To compare translation activity between wildtype and mutant UTRs, two reporter constructs of the same pair were transfected into independent wells of HEK293T cells, whereby their bioluminescence intensity was determined separately. Notably, to calibrate transfection efficiency across wells, the UTR reporter constructs were co-transfected with *Renilla* luciferase vector, and the firefly bioluminescence signal intensity was normalized to that of *Renilla*. Moreover, to control for potential variations in transcriptional activity, the relative firefly luminescence (mut/WT) of each UTR pair was further normalized to their cognate relative RNA level (mut/WT) (see Method). Our results indicate that 11 out of the 12 selected HC-sig variants displayed prominent effects on translation activity (**Fig. 3B**). Additionally, most of the tested UTR variants only altered translation efficiency but not RNA level, except for RFAXP:c.-101T>G **(Supplemental Fig. S3)**. Furthermore, we observed that 5’ UTR variants had a greater impact on translation activity relative to 3’ UTR variants (**Fig. 3C**).

**Figure 3.**
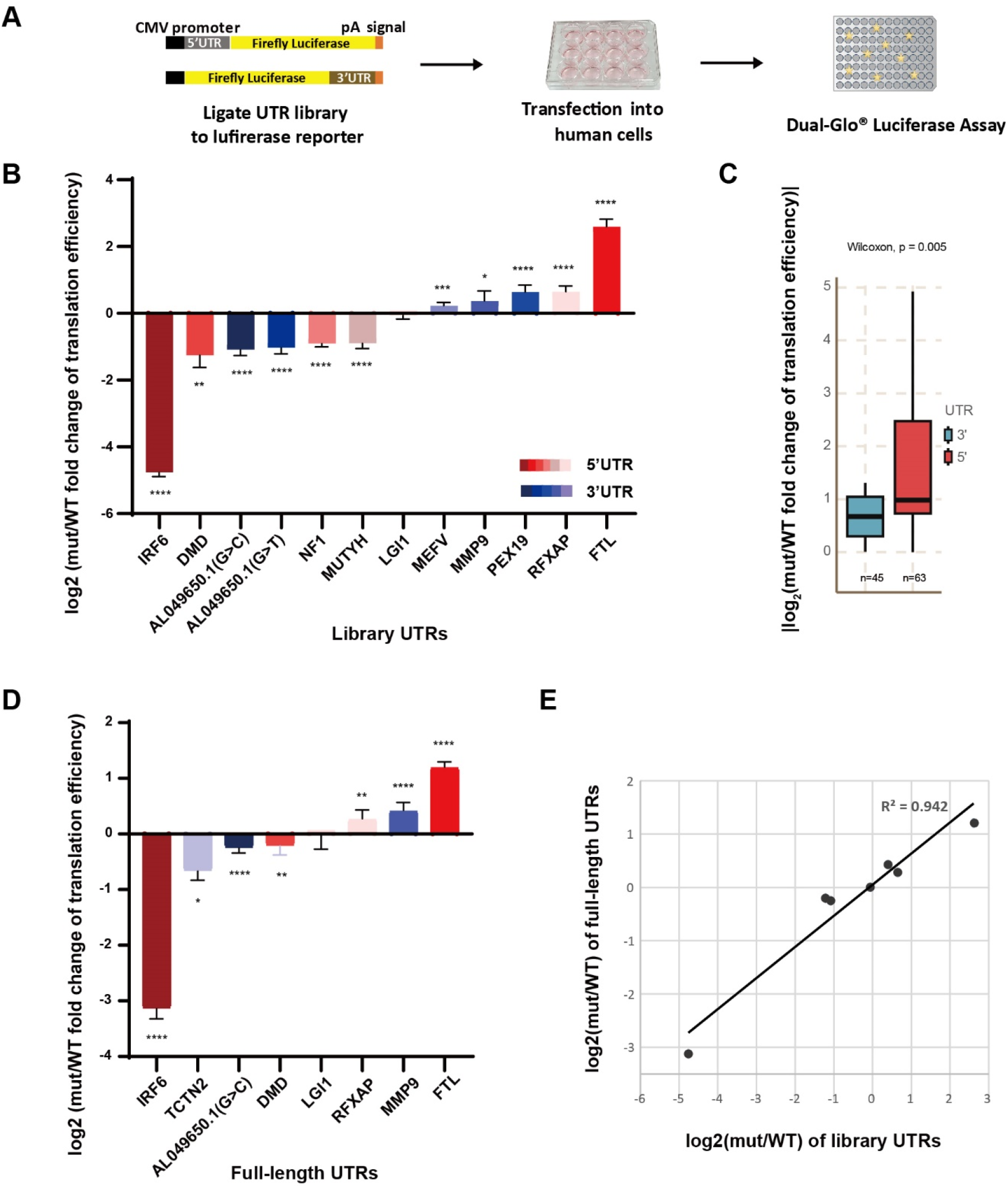
Validation of mutant effect on translation by luciferase assays. **(A)** Selected UTRs were ligated with firefly luciferase reporters and co-transfected with *Renilla* luciferase into human cells. The firefly luminescence was first normalized to the *Renilla* luminescence, and further normalized to the respective RNA expression levels to determine translation efficiency. **(B)** The log2 value of mut over WT translation efficiency was plotted ascendingly. **(C)** Absolute log2 translation efficiency changes (mut vs. WT) were compared between 5’ and 3’ UTR variants, as shown in (B). Statistical significance was assessed using a two-sided Wilcoxon rank-sum test. **(D)** Translation efficiency of eight full-length UTR pairs was examined in the same manner as in (A). UTR pairs are color-coded according to their gene origin (A&B). The individual assays were repeated three times and the differences were accessed by a two-tailed Student t-test: *: *p* < 0.05, **: *p* < 0.01, ***: *p* < 0.001, ****: *p* < 0.0001 (A&B). **(E)** Comparison of the effect of mutations on the translation efficiency of library (short) and full-length forms of UTRs.

After validating individually the variants identified in our polysome profiling assay, we proceeded to investigate their effects on translation within an endogenous genomic context, targeting a total of eight variants (five in 5’ UTRs and three in 3’ UTRs). We derived the respective full-length UTR sequences from HEK293T cells and introduced the variants into cognate UTR amplicons via site-directed mutagenesis. Next, the UTR fragments were cloned into firefly luciferase reporter plasmids, which were then used for luciferase assay. 15 hours post transfection, the translation efficiency of the full-length UTR constructs was determined and normalized as described above. Despite being embedded in a longer genomic context, 7 out of the 8 selected variants still significantly altered the translation activity of the full-length UTR constructs (**Fig. 3D**). Moreover, we observed only minor variation in the mRNA levels of the luciferase reporter across the full-length UTR constructs, indicating that the observed phenotypes could not be attributed to discrepancies in transcriptional activity or RNA stability **(Supplemental Fig. S4)**. Furthermore, UTR variant-elicited regulation was highly consistent between full-length and short-form UTRs (**Fig. 3E**), though the fold-change (mut/WT) in translation activity observed for full-length UTRs was much lower than for short-form UTRs, potentially due to the existence of additional regulatory elements in genomic milieus, which hindered or mitigated the impact of variants. Thus, our luciferase reporter assays verify the robustness of our massively parallel polysome assay results. The disparity between full-length and short-form UTRs also highlights the strength and limitation of our strategy.

### Modeling the polysome-shifting UTR variants

Recognizing that polysome-shifting variants encompass diverse classes, such as those disrupting RBP binding or secondary structure, we employed statistical learning to discern the contributions of sequence and structural features to polysome shifting. Over 3,000 features, including evolutionary conservation, folding energy, secondary structures, RBP motifs, and k-mers, were collected. Features deemed insignificant or with an area under the receiver operating characteristic (AUROC) smaller than 0.5 by univariate logistic regression were filtered out (Methods). The remaining features were further refined using elastic net regression. Twelve features were selected to elucidate polysome-shifting variants in the 5’ UTR, while eighteen were chosen for the 3’ UTR (**Fig. 4A and Supplemental Table S6)**. Notably, GAA-repeat motifs of SRSF family binding sites emerged as the most influential sequence feature for the 5’ UTR. Additionally, changes in ‘CAT’ and ‘AA’ sequences, along with the folding energy of the 5’ UTR, proved most effective in inducing polysome shifting. In contrast, for the 3’ UTR, the contribution of folding energy was less pronounced. Mutations altering GA- and T-rich sequences were identified as the strongest features promoting polysome shifting. Model accuracy was validated against a test dataset not included in the training, with both the 5’ UTR and 3’ UTR models achieving an area under the receiver operating characteristic curve (AUROC) exceeding 0.8, underscoring the model’s robustness (**Fig. 4B**). To further assess generalizability, we applied the model to an external dataset that independently measured the effects of 5’ UTR variants on polysome association . The model achieved an AUROC of 0.75 on this external benchmark (**Fig. 4C**), affirming its predictive power. Overall, our model suggest that GA- and T-rich short repeat motifs contribute to predicting mutation-induced changes in polysome association, whereas secondary structure may exert a greater influence on 5’ UTR translation regulation compared to the 3’ UTR.

**Figure 4.**
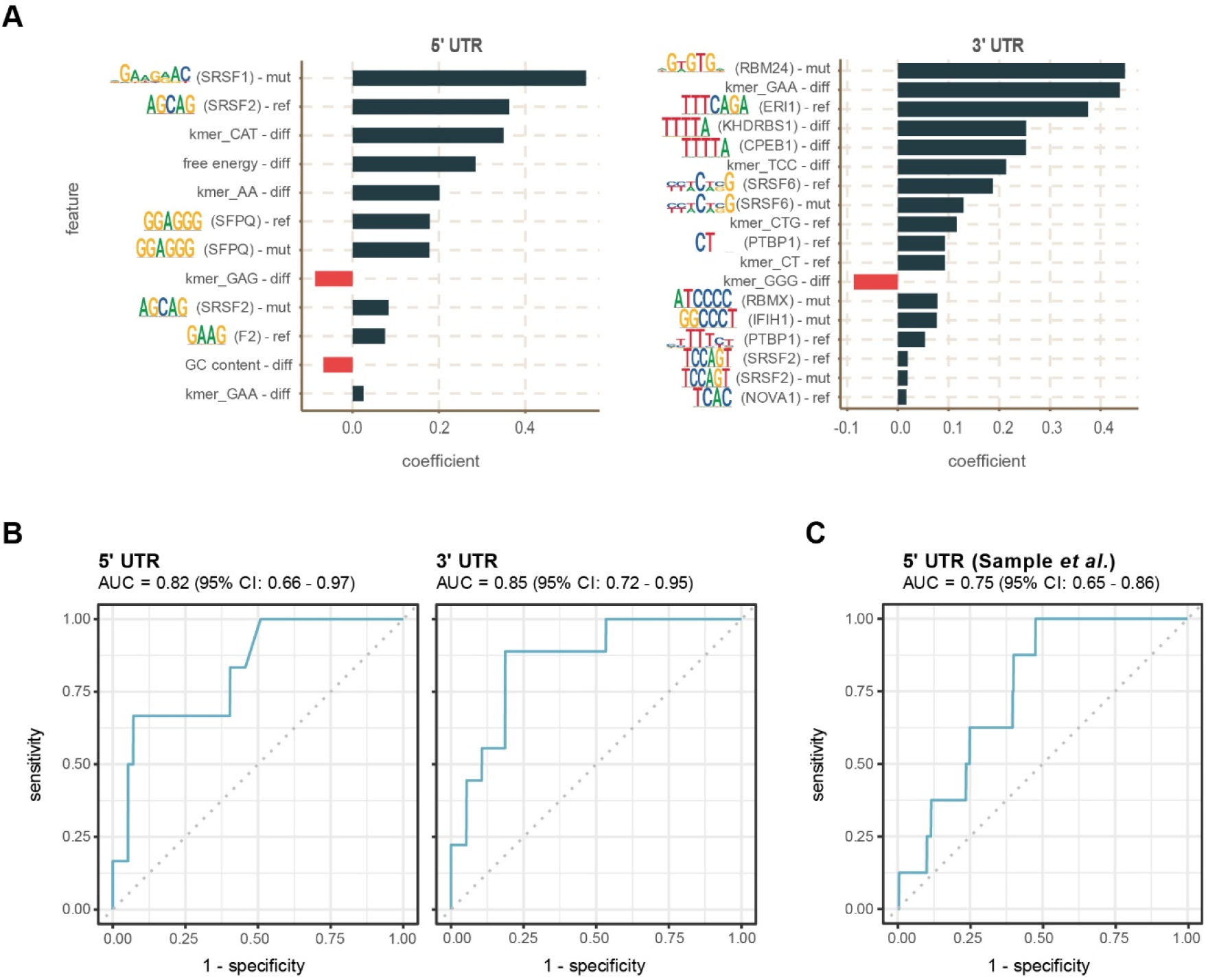
Elastic net model distinguishes polysome-shifting UTR variants. **(A)** Selected features from elastic net regression explaining the likelihood of a polysome-shifting UTR variant. A positive coefficient indicates a feature that promote polysome shifting, while a negative coefficient suppresses it. “mut” represents a feature of the mutant UTR, “ref” represents a feature of WT UTR, and “diff” indicates the difference (delta) associated with the variant. **(B)** Receiver operating characteristic (ROC) curve for the prediction of polysome-shifting variants on the test set of UTR variants. **(C)** ROC curve for the prediction of polysome-shifting variants on an external dataset of UTR variants (Sample et al., 2019a).

### Polysome-shifting UTR variants promote motif gain/loss

We hypothesized that polysome-shifting UTR variants may result in the removal or gain of functional motifs that recruit *trans* factors responsible for regulating translation. Accordingly, we postulated that polysome-shifting variants would display distinct sequence properties. Therefore, first we investigated the nucleotide composition of polysome-shifting variants. We found that variations in C nucleotides (including gain and loss) in 5’ UTRs were relatively inert in terms of translational alteration, whereas that was not the case for C in 3’ UTRs **(Supplemental Fig. S5A and Table S7)**. Subsequently, we investigated the RBP-binding motifs of polysome-shifting UTR variants. Our results show that GAAGAA and T-rich motifs are enriched in HC-sig 5’ UTRs, whereas G-rich and YTTC motifs are enriched in HC-sig 3’ UTRs **(Supplemental Fig. S5B and Table S7,** Y represents T or C). Furthermore, recognizing the intricate relationships between binding motifs and their corresponding RBPs, we conducted an additional analysis focusing on the RBPs themselves. This analysis suggested that TRA2B and members of the HNRNP family can bind HC-sig UTRs to regulate their translation **(Supplemental Fig. S5C)**. Extending this analysis, we calculated if changes in RBP-binding motifs are associated with changes in polysome profiles, which revealed that changes of SRSF1/2-, PABPN1-(AGAAG motifs) and CELF2-(T-rich motifs) binding sites in 5’ UTRs frequently caused polysome shifting, so did mutations of binding sites for KHSRP (AGGG motifs) in 3’ UTRs (**Fig. 5A**). These findings were consistent with the elastic net model, in which GAA-repeat motifs recognized by SRSF-family proteins emerged as prominent features associated with polysome-shifting UTR variants.

**Figure 5.**
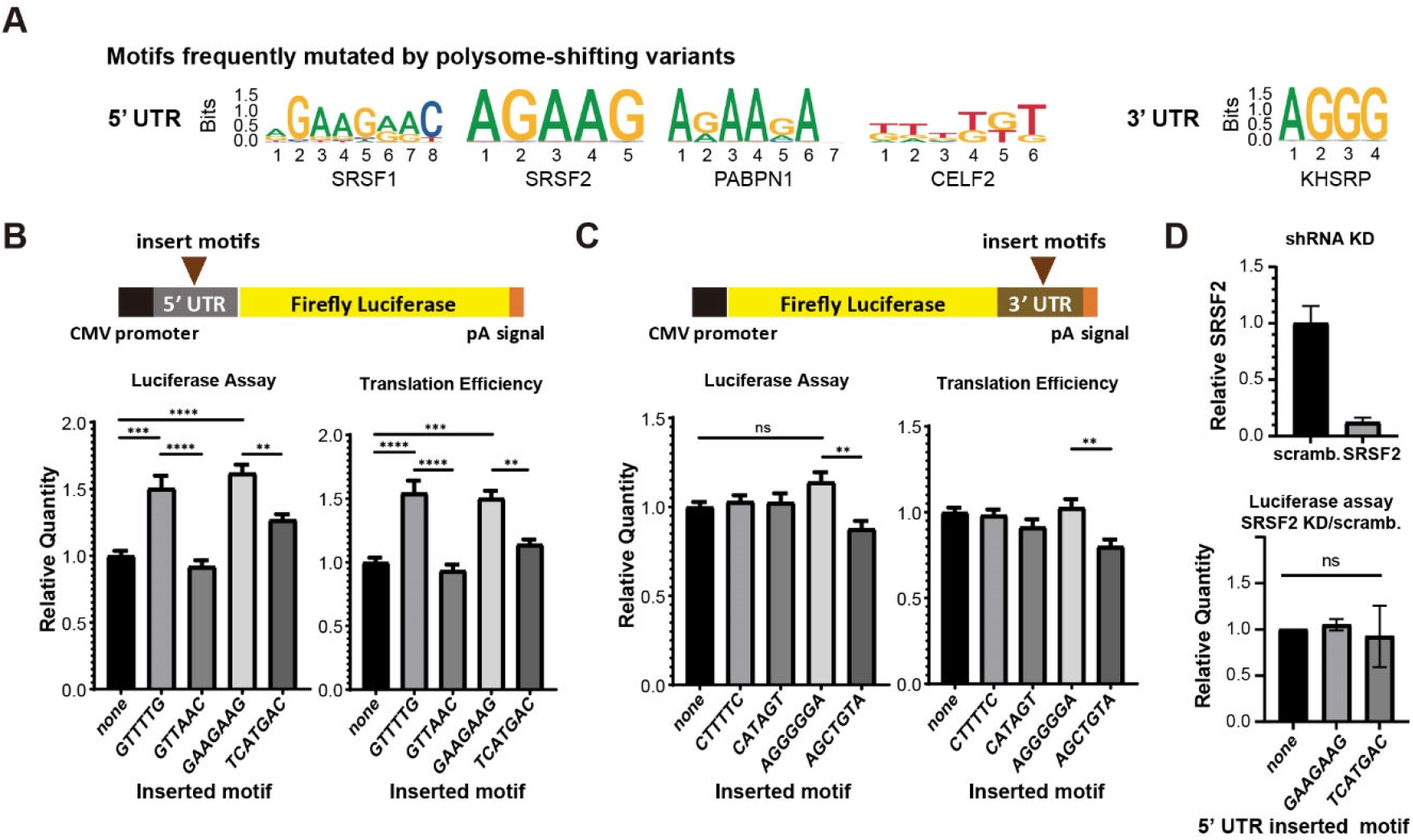
Identification of polysome-shifting RBP-binding motifs. **(A)** Motifs significantly more frequently mutated by HC-sig variants. **(B)** Two 5’ UTR-enriched motifs from (A), i.e., GTTTTG and GAAGAAG, were inserted into a 5’ UTR of firefly luciferase to determine their luciferase activity and normalized translation efficiency. Mutant forms of these two motifs, GTTAAC and TCATGAC, were generated for comparison. **(C)** Similar to the 5’ UTR design in (B), the 3’ UTR-enriched motifs CTTTTC and AGGGGGA were examined against their mutant forms, i.e., CATAGT and AGCTGTA. The individual assays were repeated four times. Statistical significance was determined by a two-tailed Student t-test: **: *p* < 0.01, ***: *p* < 0.001, ****: *p* < 0.0001. **(D)** Upper: Relative SRSF2 transcript levels, as determined by RT-qPCR. Lower: Relative luciferase activity with various 5’ UTR motifs, normalized to scrambled shRNA infection.

To further validate the RBP-binding motifs identified by our enrichment analysis, we examined the effect of selected motifs on reporter expression by inserting them into a segment of the *MUTYH* 5’ UTR or *MEFV* 3’ UTR that contains suitable cloning sites (see Methods) and then perturbing its effect by mutating it so that it retained the same length but rendered it non-RBP-binding according to the ATtRACT RBP motif database (Giudice et al., 2016). For 5’ UTR, GTTTTG and GAAGAAG motifs were selected for the test. Both selected motifs significantly increased the translation efficiency of luciferase reporters without affecting RNA expression level (**Fig. 5B**). For 3’ UTR CTTTTC and AGGGGGA motifs were selected for the test, yet neither of them significantly altered reporter expression. However, mutation of the AGGGGGA motif into AGCTGTA significantly reduced translation efficiency, indicative of a context-dependent effect of RBP-binding motifs (**Fig. 5C**). This observation is consistent with additional findings where variants that create or disrupt specific RBP binding sites—such as SRSF1/2 (e.g., in *DMD* and *NF1*; **Fig. 3 and Supplemental Fig. S4)** and KHSRP (e.g., in *AL049650.1*; **Fig. 3 and Supplemental Figs. S4 & S5)**—led to significant changes in translation efficiency within their native UTR contexts.

Additionally, to assess the impact of RBPs on translation regulation through the short repeat sequence, we conducted a knockdown of SRSF2 using shRNA. This RBP was chosen due to its cytoplasmic localization (Thul et al., 2017) and abundant expression in HEK293T cells compared to other candidate RBPs. The results revealed that depleting SRSF2 eliminated the translation enhancement mediated by GAAGAAG, while having no effect on a non-SRSF2-binding motif (**Fig. 5D**). While SRSF2 has not been directly implicated in translation regulation, some members of the SRSF family have been reported to function as translation regulators (Sliskovic et al., 2022). Notably, a mutation in the *NUMA1* 5’ UTR that disrupted SRSF9 binding was previously demonstrated to decrease translation efficiency (Lim et al., 2021), providing further support for the involvement of SRSF family proteins in translation regulation. Together, these results support the existence of mutational hotspots on short repeated sequences for polysome-shifting UTR variants and a collaborative regulatory effect of binding motifs.

### Polysome-shifting mutations alter RNA secondary structure

Given that RNA secondary structure is believed to control translatability in addition to primary sequences (Georgakopoulos-Soares et al., 2022; Leppek et al., 2018), we determined the changes in folding energy elicited by sequence variations. To do so, we compared the folding energy change for variants in the HC-sig and HC-nonsig groups, which revealed that the 5’ UTR HC-sig variants significantly weakened the folding structures and tended to prompt greater structural change than HC-nonsig variants (**Fig. 6A**). Furthermore, the HC-sig WT exhibited robust folding energy, potentially underlying their susceptibility of folding perturbations. In the same vein, RNA secondary structure predictions showed notable structural changes caused by 5’ UTR single-nucleotide variations. As an example of the latter, we present the case of a 5’ UTR single nucleotide variation in C-C chemokine receptor type 5 (CCR5) in Figure 6B. These results suggest that structural alterations of 5’ UTRs by sequence variation could be a potential causal factor of the translation defects.

**Figure 6.**
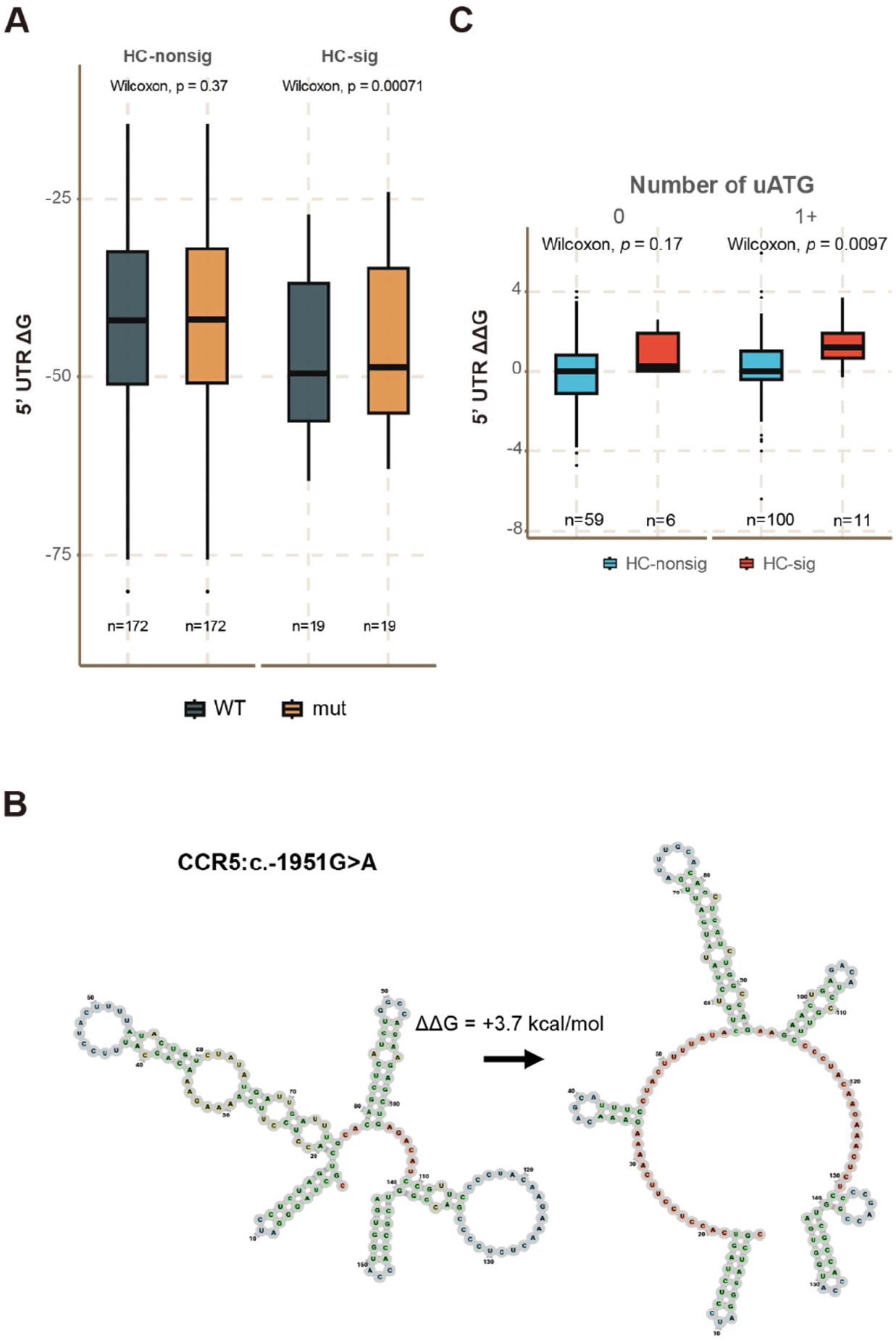
5’ UTR Polysome-shifting mutations elicit structural changes. **(A)** Significantly weaker folding energy of 5’ UTR polysome-shifting variants. **(B)** Structural change induced by a CCR5 5’ UTR single nucleotide change, as determined by RNAfold. **(C)** Significant differences in ΔΔG values for 5’ UTRs that contain upstream ATG (uATG).

Since it has been shown previously that the usage of an uATG is affected by the surrounding UTR structure and can regulate translation of primary ORFs (Guenther et al., 2018; Xiang et al., 2023), we examined if the significant structural change elicited by HC-sig variants could be associated with the presence of uATG. Indeed, by contrasting UTR variants that did not alter uATG counts, we found that uATG-hosting UTRs were strongly associated with structural changes (**Fig. 6C**). Therefore, while changes in uATG may not be common explanatory factors for polysome-shifting mutations, our results suggest that structure-modifying UTR variants may control primary ORF translation partly by interfering with translation initiation from a uORF.

### Modeling the polysome occupancy of 5’ UTRs

Because 5’ UTRs contain multiple features, including RBP-binding motifs and RNA folding energy, that may jointly influence translation, we next asked whether these features could explain variation in polysome occupancy. We performed high-resolution polysome profiling of the UTR library, separating transcripts into free RNA (Free), 40S, monosome (1 ribosome), and polysome fractions containing 2-8 ribosomes (**Fig. 7A**). We evaluated several metrics of ribosome association and found that the polysome fraction normalized to the free and 40S fractions [Poly/(Free + 40S)] showed the strongest agreement with the change in translational efficiency between WT and mutant pairs measured using luciferase reporters (Spearman r = 0.744) (**Figs. 3 and 7B**). We therefore used this index as a quantitative measure of polysome occupancy and assigned the direction of occupancy change (increased or decreased) for each polysome-shifting variant **(Supplemental Table S4)**.

**Figure 7.**
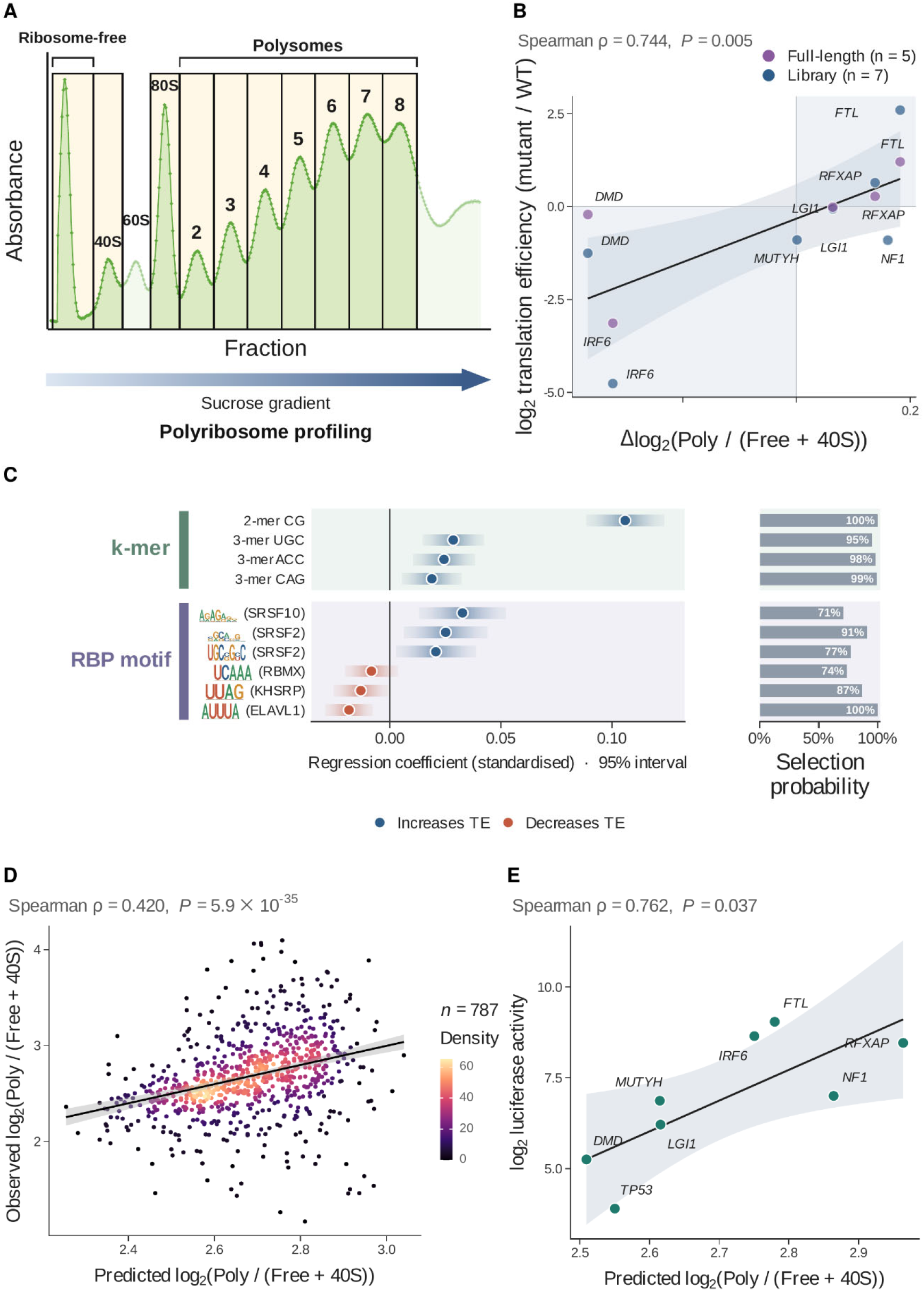
A 5’ UTR sequence model predicts translation efficiency and agrees with luciferase assays. (**A**) Schematic of polysome profiling. Translation efficiency (TE) was defined as log₂[polysome (F2 to F8) / (free + 40S)] and averaged over three biological replicates. (**B**) Mutation effects (mut vs. WT) measured by the MPRA (ΔTE) and by luciferase assays (log₂ TE ratio, determined as in Figure 3). Blue, library constructs (*n* = 7); purple, full-length constructs (*n* = 5). Shaded quadrants indicate agreement in direction. **(C)** Features retained by stability selection (selection probability > 0.7 across 100 random subsamples of half the data, right bars) and their standardized coefficients with 95% confidence intervals from weighted linear regression. Blue, positive; red, negative. RBP motifs are shown as ATtRACT sequence logos. (**D**) Observed versus predicted TE in the held-out test set (*n* = 787). (**E**) Predicted TE versus luciferase activity (firefly normalized to *Renilla*) of eight wild-type UTRs. Values are means of two biological replicates, each with three technical replicates. Lines in (B), (D) and (E) are linear fits with 95% confidence intervals. Spearman ρ and *P* values are shown above each panel.

To identify 5’ UTR features associated with polysome occupancy, we first selected representative features from highly correlated feature groups (for example, CG dinucleotide frequency and overall GC content) and then applied standardized multivariable regression to estimate their relative effects, statistical significance, and confidence intervals (see Methods). Of the 2,850 WT 5’ UTRs in the library, 2,362 passed the read-depth and quality-control criteria and were included in the model. The resulting model revealed positive associations of GC-rich k-mers with polysome occupancy. Notably, GC-rich sequence features were also associated with RNA stability (Courel et al., 2019; Su et al., 2025), suggesting a potential link between UTR-mediated RNA stability and translational efficiency. GA-repeat motifs recognized by SRSF-family proteins were also positively associated with polysome occupancy (**Fig. 7C**), consistent with their enrichment among polysome-shifting UTR variants identified independently in our variant analysis (**Figs. 4 and 5**).

The multivariable model significantly explained variation in the observed polysome occupancy (**Fig. 7D, Supplemental Table S4)**. Moreover, model-derived polysome occupancy highly correlated with luciferase activity measured independently across eight WT 5’ UTRs (Spearman r = 0.762) (**Fig. 7E**), providing functional support for the relationships captured by the model. Together, these results identify sequence and regulatory features associated with 5’ UTR-dependent ribosome loading and provide a quantitative framework linking UTR composition and RBP-binding motifs to translational output.

### The translation defect of a uORF-creating variant can be rescued by RNA editing

While uATG in the 5’ UTR was not identified by the model as a widespread feature explaining polysome shifting, we hypothesize that it may still be a crucial factor. Its significance, however, might not have been universally captured across our library. Genome-wide analysis clearly indicated an enrichment of uORF disruption in the pathogenic UTR mutations (**Fig. 1B**). Therefore, we aimed to dissect the regulatory mechanisms of uORFs underlying the effects of polysome-shifting variants and develop rescue strategies against pathogenic uORF mutations.

First, we examined whether the presence of uATG in general leads to lower translation in reporter assays. We observed that the presence of uATG in 5’ UTRs of firefly luciferase was correlated with lower protein outputs (**Fig. 8A, Supplemental Table S8)**. Furthermore, two of the validated UTRs—Ferritin light chain (FTL) and Interferon Regulatory Factor 6 (IRF6)—demonstrated extreme translation perturbations (**Fig. 3**). In a study by Luscieti et al., an FTL variant (c.-161C>T) was characterized as preventing 5’ UTR stem-loop formation (Luscieti et al., 2013). As a consequence, the recruitment of suppressive iron regulatory proteins mediated by the 5’ UTR stem-loop is disrupted, leading to an increase in translation activity. An IRF6 variant (c. −4609 G>A) was found previously to be associated with the developmental disorder Van der Woude syndrome (de Lima et al., 2009). The G>A variant was predicted to generate an out-of-frame uORF, but no experimental evidence has been provided yet. As shown in Figure 3, the corresponding full-length mutant UTR displayed a dramatic 9-fold reduction in translation efficiency, strongly indicating that presence of the uORF represses primary ORF translation. To verify the role of uATG, we performed western blotting to measure protein levels directly and determine protein size. To do so, we generated a third reporter construct that harbored the uORF in-frame to the coding region by deleting one nucleotide (**Fig. 8B**). When probed with a luciferase-targeting antibody, the signal intensity for the out-of-frame mutant after normalization to the wildtype was only 0.089 (**Fig. 8C**). This suggests that only a minute fraction of ribosomes may reinitiate at the canonical site or undergo leaky scanning at the uATG. Notably, the in-frame mutant, which represents a combination of the extended and the original forms, displayed a 0.24-fold signal intensity, indicating that uATG, despite lower initiation efficiency, markedly suppresses initiation at the canonical site. Compared to wildtype, we determined 0.6-fold translation efficiency, as measured by luciferase reporter assays, upon overexpression of the in-frame mutant (**Fig. 8D**) and no significant change in mRNA level was observed **(Supplemental Fig. S4)**. The discrepancy between protein abundance and luciferase intensity indicates that the extended form of the luciferase reporter may contribute to higher luminescence intensity. Together, these results support a role for the variant-created uATG in repressing translation of the downstream canonical ORF, independently of changes in mRNA abundance.

**Figure 8.**
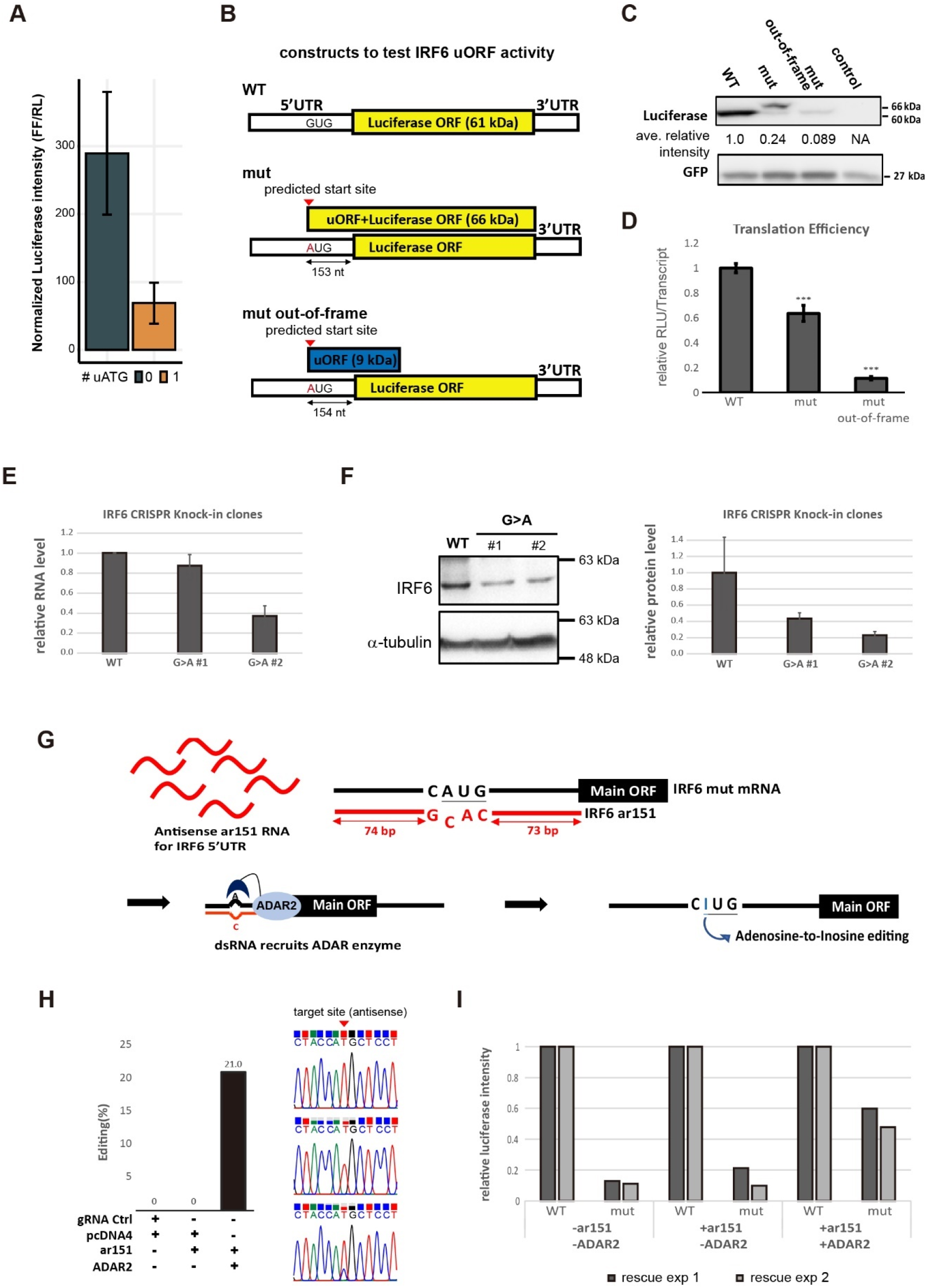
5’ UTR mutations cause translation inhibition by generating upstream ATG. **(A)** Luciferase activity of eight UTRs with or without a uATG in the 5’ UTR. Error bars represent standard errors. **(B)** Constructs of the mutant IRF6 5’ UTRs. Mutant (mut) IRF6 generates an in-frame luciferase protein from an upstream ATG, whereas the uORF and luciferase are coded in different frames for the out-of-frame mutant (mut out-of-frame). **(C)** Western blots showing a reduction in luciferase protein levels following 5’ UTR mutations. **(D)** Luciferase assays showing a reduction in translation efficiency due to 5’ UTR mutations. RLU: relative luminescence units. The individual assays were repeated three times. Statistical significance was determined by a two-tailed Student t-test against WT: ***: *p* < 0.001. **(E)** Relative expression of IRF6 RNA in CRISPR knock-in HEK293T cells. The G>A clones carry the IRF6:c.-4609G>A variant. IRF6 RNA expression was assessed by four independent RT-qPCR experiments and normalized to GAPDH. The error bars represent standard errors. **(F)** IRF6 protein expression in CRISPR knock-in clones. The bar graphs summarize three independent Western blot analyses of IRF6, with normalization to α-tubulin protein levels. The error bars represent standard errors. **(G)** Design of guided ADAR-mediated editing to remove a uORF. **(H)** Editing efficiency achieved by guided ADAR, as determined by Sanger sequencing. **(I)** Restoration of translation efficiency by means of guided ADAR editing, as determined by luciferase assays.

To further validate the endogenous effect of the novel upstream ATG (uATG), we generated CRISPR knock-in clones carrying the IRF6:c.-4609G>A variant and examined its impact on gene expression. The introduction of the uATG reduced RNA levels to 88% and 37% of the wild-type in two independent clones (**Fig. 8E**), and protein levels to 44% and 23%, respectively (**Fig. 8F**), resulting in an overall reduction of translation efficiency to 50–62%. Although the uORF-mediated impairment of translation efficiency in IRF6 was clearly validated, we were unable to detect the corresponding micropeptide in the out-of-frame constructs (data not shown). It remains unclear if micropeptides, if translated, play any role in pathogenesis. However, patients who have gained a uORF in the IRF6 5’ UTR phenocopy Van der Woude patients hosting loss-of-function IRF6 mutations (de Lima et al., 2009).

Lastly, we aimed to develop rescue strategies that may be useful for future therapeutic purposes. We selected site-directed RNA editing using adenosine deaminase acting on RNA (ADAR) enzyme for this purpose to avoid genome editing and potential off-target effects. Upon binding of an antisense RNA to a target transcript, the ADAR enzyme catalyzes adenosine-to-inosine editing. Inosine is subsequently interpreted as guanosine during translation (Merkle et al., 2019; Qu et al., 2019). To achieve high editing efficiency, we overexpressed the antisense RNA coding construct and ADAR2 protein (**Fig. 8G**). The transcribed antisense RNA encompasses a T>C mismatch that signals the complementary nucleotide for editing. First, we assessed editing efficiency by means of Sanger sequencing (**Fig. 8H**). Only co-expression of antisense RNA and ADAR2 resulted in detectable editing, with an efficiency of ∼21%. Luciferase signal intensities were also validated (**Fig. 8I**). Consistently, expression of the antisense RNA alone had a very limited effect, but co-introduction of ADAR2 significantly increased the protein expression to ∼50% the level of wildtype. Our promising results advocate the potential of clinical use of guided ADAR-RNA editing to reverse uORF-mediated translation suppression and thus alleviate the impact of the disease.

## DISCUSSION

In this study, we applied massively parallel polysome assays to examine the effect of mutations in 5’ and 3’ UTRs in terms of RNA translatability (**Fig. 2**). Our results show that the effect we observed in our high-throughput profiling is reliable, though it could be diluted somewhat in a more complex genomic context (**Fig. 3**). We have shown that T- and GAA-repeat motifs are sensitive to translation-altering mutations (**Fig. 4 and 5**), underscoring the involvement of both *cis*-regulatory sequences and *trans*-acting RBPs in regulating RNA translatability. Additionally, we identified the secondary structure of 5’ UTRs as a critical factor controlling RNA translatability (**Fig. 6**). We observed that alterations in folding energy, especially in uORF-hosting 5’ UTRs, can have detrimental effects, suggesting structure-mediated uORF translation. These findings were summarized by regression models and robustly explained the polysome-shifting UTR variants and the UTR polysome occupancy (**Fig. 4 and 7**). Additionally, the creation of uORFs in 5’ UTRs through mutation adversely impacts primary ORF expression (**Fig. 8B-F)**, consistent with our finding of pathogenic mutant hotspot at uORF translation start sites (**Fig. 1B**). We also reveal that ADAR-mediated editing can be deployed to rescue primary ORF translation (**Fig. 8G-I)**. Our study emphasizes the significant role of precise UTR translation regulation in gene expression and offers a molecular explanation for the impact of UTR mutations on pathogenicity. Below, we discuss some of the strengths and limitations of our study.

### Massively parallel evaluation of translation efficiency

Translation efficiency is a key post-transcriptional regulation to control gene output. Efforts have been made to assess translation efficiency systematically in terms of gene structures and sequences (Jia et al., 2020; Leppek et al., 2022; Niederer et al., 2022; Oikonomou et al., 2014; Sample et al., 2019a; Savinov et al., 2021; Slutskin et al., 2018; Wissink et al., 2016; Zhao et al., 2014). One way to measure translation efficiency is according to protein output. For instance, in MPRA where reporter sequences are fixed and fused with test sequences, the expression of the reporter fluorescent proteins is determined by Fluorescence Activated Cell Sorting (FACS), with expression being normalized to the level of the other fluorescent reporter from the same construct to account for experimental bias. The tested sequences within each bin are then determined by sequencing. Mass spectrometry can be employed to determine quantitatively endogenous protein levels (Baek et al., 2008). Although this method is relatively quantitative, MPRA has the leverage over RNA stability, especially if the reporter is fused with various UTRs. Moreover, the results for endogenous proteins are influenced by protein turnover, which itself is a very important means of controlling gene expression. Thus, although protein output is a crucial readout of gene expression, this approach does not provide a direct measurement of translation efficiency.

The alternative method to measure translation is through polysome profiling. For endogenous genes, ribosome footprinting can be employed to quantify transcript fragments associated with ribosomes. For MPRA, in which the coding footprint is identical for the whole library, polysome profiling is used instead. Rather than sequencing and then counting transcript fragments attached to ribosomes, polysome profiling does not shear off the transcript, but only fractionates RNAs according to the number (size) of associated ribosomes through ultracentrifugation in a sucrose gradient, representing the most direct way to measure the efficiency of RNAs to recruit ribosomes. However, there are some limitations to this method. First, there is considerable variation in how polysome fractions are used. Some studies have used the average number of ribosomes on each reporter, with each fraction (including the ‘ribosome-free’ fraction) being considered and the average ribosome number being calculated according to the relative distribution of the transcript in polysomes (Leppek et al., 2022; Sample et al., 2019a, 2019b). In contrast, others use only some of the fractions to infer translation efficiency. For example, percentage translated RNAs in polysomes can be used to infer ribosome loading (Jia et al., 2020; Schuster et al., 2023), and the monosome to input ratio (recruitment score) can be calculated to infer ribosome recruitment efficiency (Niederer et al., 2022). These alternative approaches to analyzing the data create perplexity to interpret polysome profiles. Another major limitation of polysome profiling is its dependency on promoters. It has been reported that whether polysome profiles reflect translation efficiency is contingent on the promoters driving gene expression (Kong et al., 2008). It remains unclear if the strong expression promoters (such as the CMV promoter) commonly used for MPRA are optimal for revealing differences in polysome profiles. Moreover, because not all ribosomes translate actively on transcripts, they can be slowed, stalled, or collided on a transcript (Meydan & Guydosh, 2021). Although it has been proposed that the major bottleneck of the translation reaction is translation initiation, elongation also contributes to translation efficiency (Riba et al., 2019). Thus, whether the number of ribosomes associated with a transcript truly reflects translation efficiency, especially for endogenous transcripts in diverse coding contexts, remains elusive.

### Severe translation defects caused by 5’ UTR point mutations

The 5’ UTR serves as the major ribosome entry site for translation. In our data, we observed remarkable changes in translation efficiency elicited by 5’ UTR point mutations (**Fig. 3**), with IRF6:c.-4609G>A and FTL:c.-161C>T protein outputs being reduced 10-30-fold without affecting respective RNA levels **(Supplemental Figs. S3 and S4)**. The impacts of 3’ UTR mutations appear to be more additive and context-dependent, consistent with its nature of greater length variation. The most severe UTR mutation we identified in our MPRA dataset is IRF6:c.-4609G>A, which has been observed in Van der Woude syndrome patients. This 5’ UTR mutation has been suggested to suppress IRF6 translation by generating a uORF. We have demonstrated herein that IRF6:c.-4609G>A creates a strong upstream translation start codon, dramatically reducing protein output from the primary ORF (**Fig. 8B-F)**. We reasoned that a G>A mutation represented an optimal target for ADAR editing, allowing us to switch A back to I, which is read as G by the cellular machinery (**Fig. 8G-I)**. Another advantage of ADAR is that editing operates at the RNA level within the potential treatment window, thereby restoring desired protein function without causing any permanent alterations to the DNA.

Ferritin is composed of L (FTL) and H (FTH) subunits, and it is responsible for regulating the storage and intracellular distribution of iron. FTL:c.-161C, one of the UTR variants we observed as eliciting strongly perturbed translation, has been mapped as part of a noncanonical upstream start codon (CAG, TISdb) (Wan & Qian, 2014). It is likely that the C>T mutation disrupts uORF initiation, thereby releasing repression of the primary ORF, and promoting the ∼10-fold increase in translation. However, the 5’ UTR of FTL is also responsible for forming an IRE stem-loop (Klausner et al., 1993). The FTL:c.-161C>T mutation likely disrupts this hexanucleotide loop structure, preventing it from interacting with the Iron Regulatory Proteins IRP1 and IRP2 (Luscieti et al., 2013). Since IRPs bind to IREs under iron-deficient conditions to suppress translation of both ferritin subunits (Muckenthaler et al., 1998), mutations that abolish IRP-IRE interactions elicit aberrant production of ferritins and are the major causative mutations of Hereditary Hyperferritinaemia Cataract Syndrome. Thus, our massively parallel polysome profiling has uncovered known and novel mutations that cause translation defects, serving as a genetic diagnostic tool for identifying crucial and clinically relevant regulatory mutations.

## DATA AVAILABILITY

All raw and processed sequencing data generated in this study have been submitted to the NCBI Gene Expression Omnibus (GEO; https://www.ncbi.nlm.nih.gov/geo/) under accession number GSE229492 (reviewer token oxcpuuechhihzuf) and GSE349290 (reviewer token gxmvyksepriphar).

Codes used for the analyses in this study have been deposited at https://github.com/chienlinglin/modeling-UTR-variants-polysome/

Other databases used in the study:

AREsite2: http://nibiru.tbi.univie.ac.at/AREsite2

ATtRACT: https://attract.cnic.es/download

## ACKNOWLEDGEMENTS

We thank the Genomics Core and the Bioinformatics Core of the Institute of Molecular Biology (IMB), Academia Sinica, for performing the amplicon sequencing and for providing computing resources. We thank all members of IMB for tremendous help and support.

## AUTHOR CONTRIBUTIONS

C-L L and H-L C designed the experiments. W-P L, Y-C C and C-H Y established the MPRA. J-Y S, Y-L W and M K carried out modeling and analyses. W-P L, Y-C C, H-L C, Y-H C, M L, Y-L K, B-J C, Y-T Hsieh and C-H Y performed experiments. Y-T Huang supervised the statistical analysis. J-Y S, W-P L, Y-C C, Y-L W and C-L L wrote the manuscript.

