## Supplemental Figures for "Massively Parallel Polyribosome Profiling Reveals Translation Defects of Human Disease-Relevant UTR Mutations"

### Figure S1

**5'UTR**

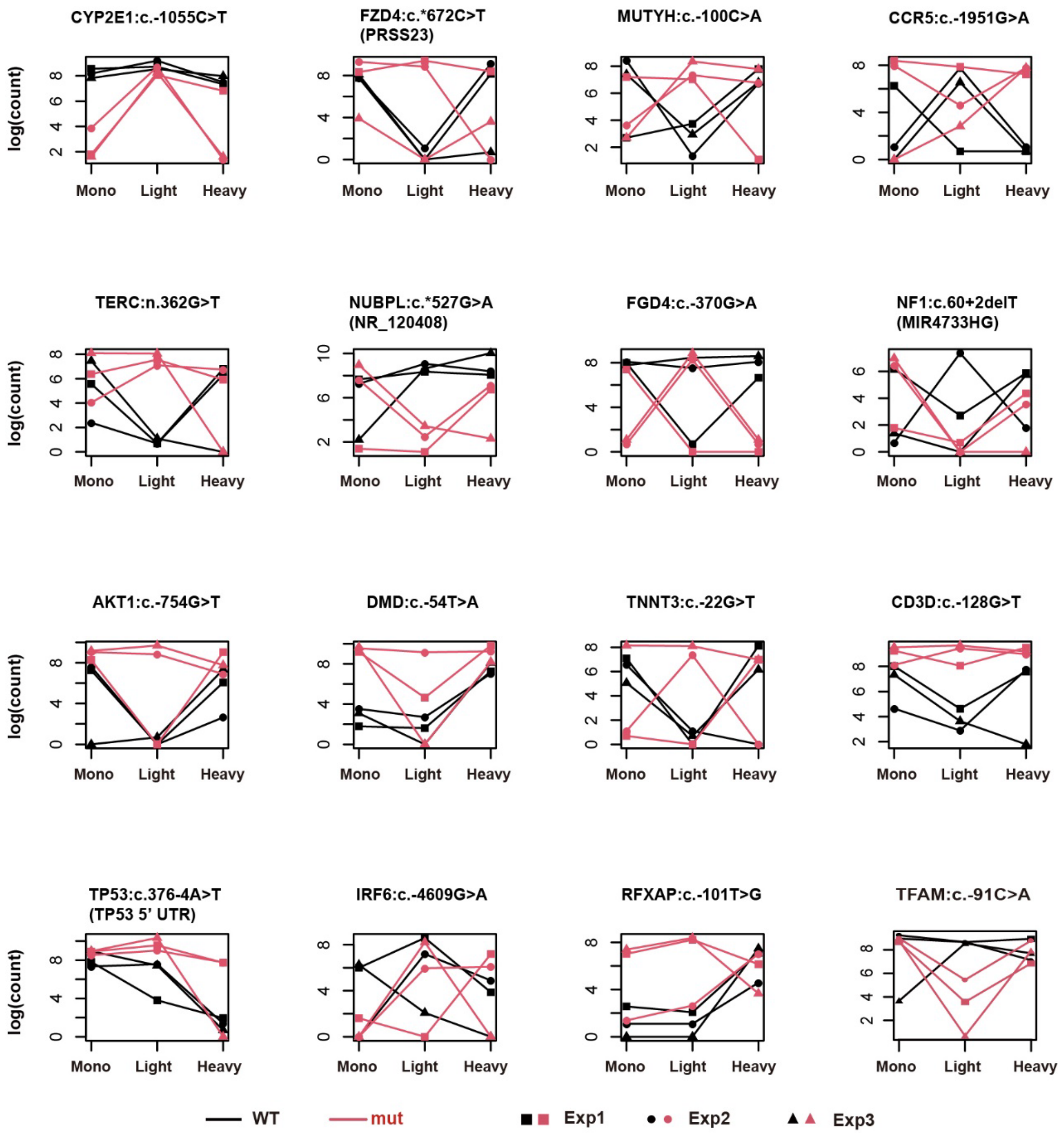


**Figure S1. 5' UTR pairs that significantly differed in polysome distribution, related to Figure 2B.** For variants that are mapped to multiple gene regions, the alignment reported in the disease databases is presented on top, with the alternative alignment indicated in parentheses. Certain noncoding genes have been proved to harbor coding potential, and therefore their regulatory sequences were incorporated into the study.

**Figure S2**

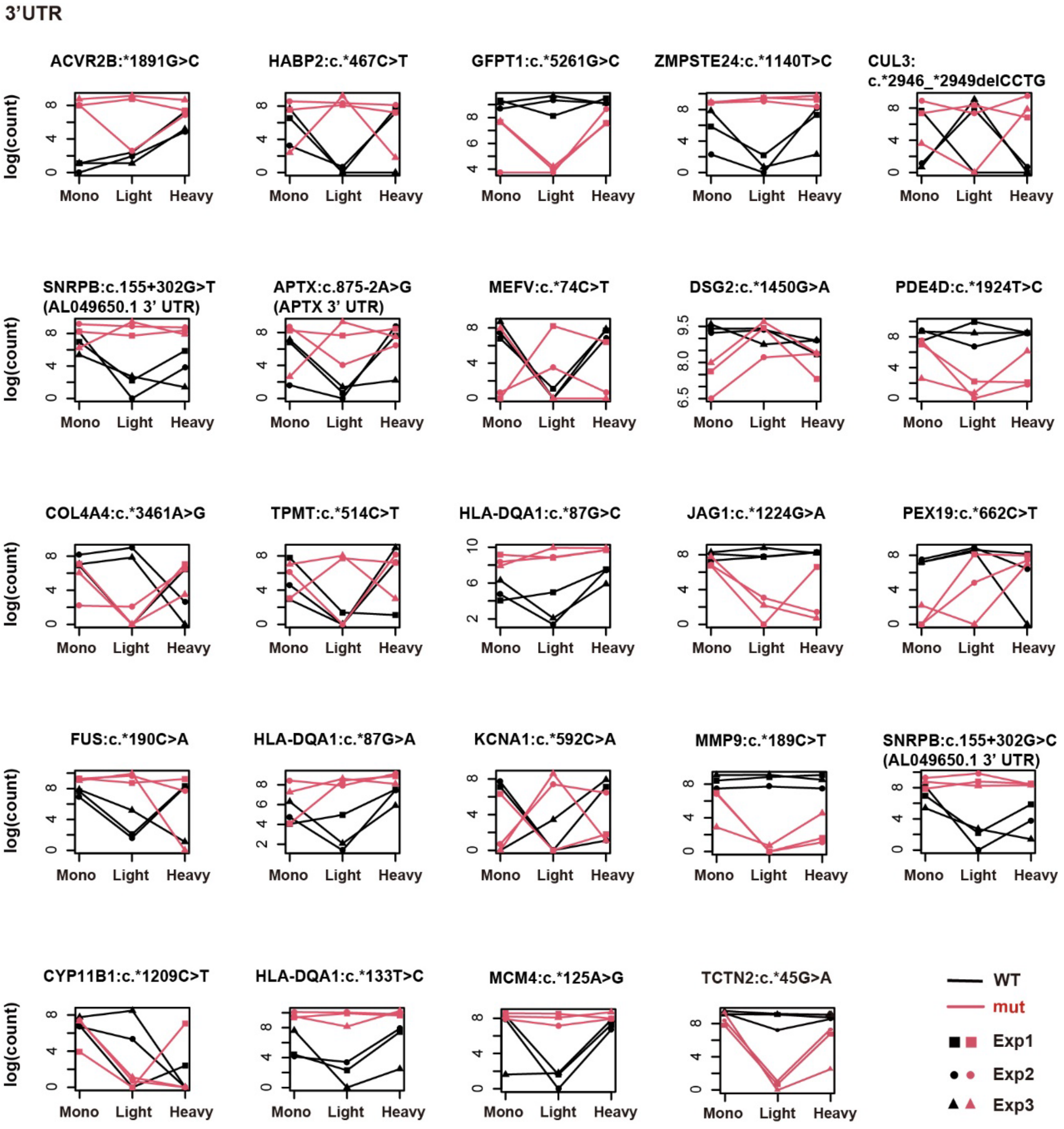


**Figure S2. 3' UTR pairs that significantly differed in polysome distribution, related to Figure 2B.** For variants that are mapped to multiple gene regions, the alignment reported in the disease databases is presented on top, with the alternative alignment indicated in parentheses.

**Figure S3**

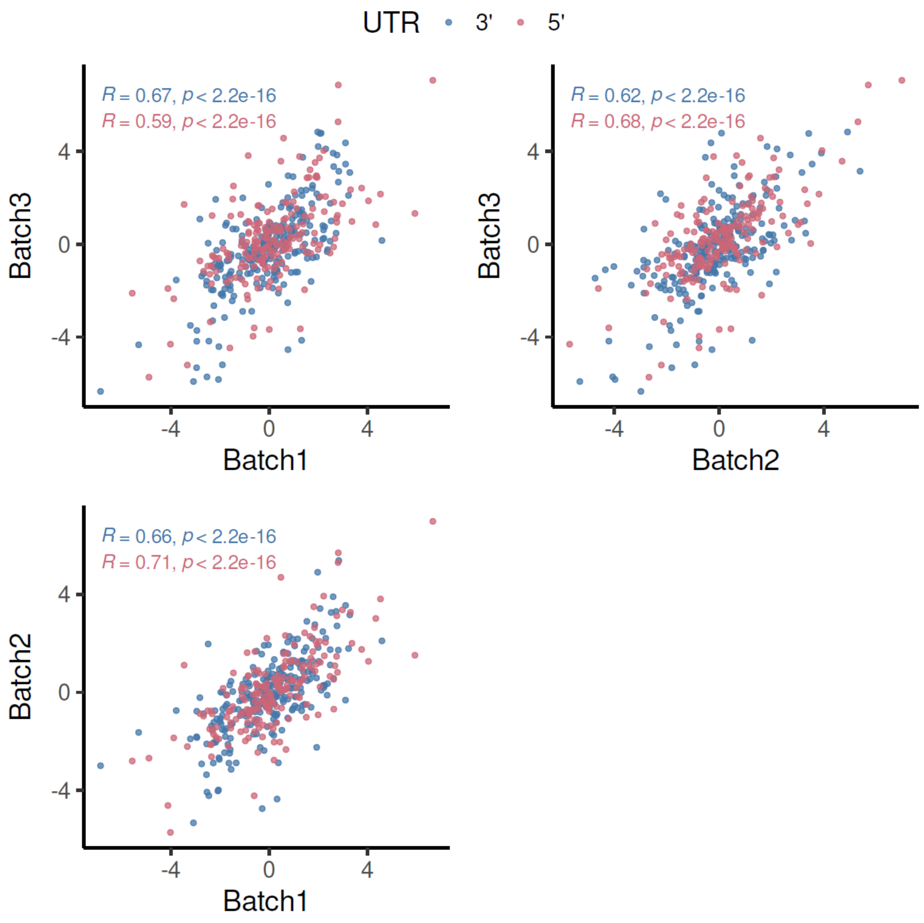


**Figure S3. Data reproducibility of the massively parallel polysome profiling.** Scatterplots show the Pearson correlation of log-transformed relative mutant-to-wild-type, $\ln(\frac{\text{mut}}{\text{wt}})$, transcript ratios across three independent experimental batches for the high-confidence dataset, calculated from the sum of monosome, light, and heavy polysome fractions. Correlation coefficients and P values are determined by the Pearson correlation test. Red dots represent 5′ UTR library constructs, and blue dots represent 3′ UTR library constructs.

**Figure S4**

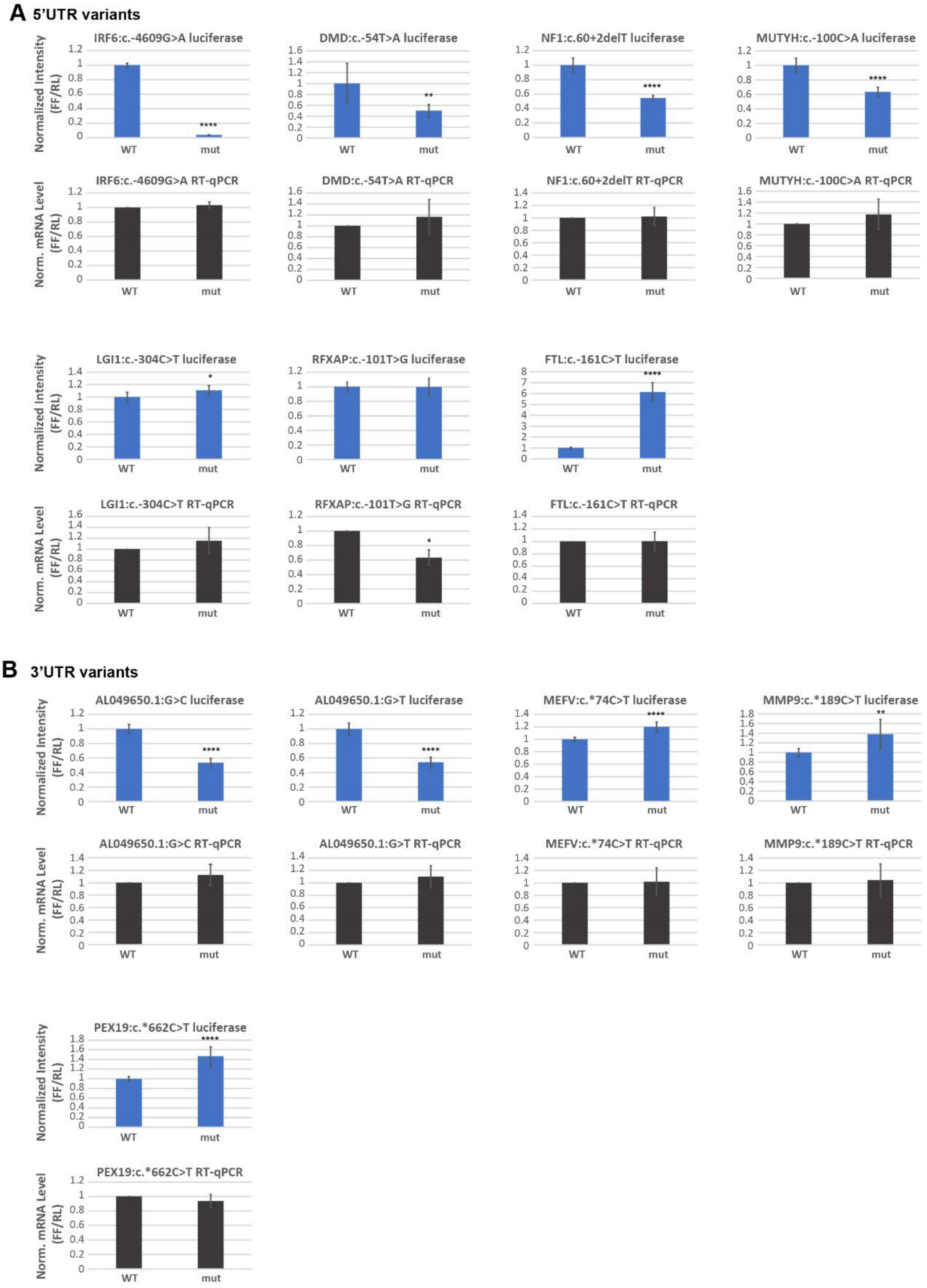


**Figure S4. Validation by luciferase assays of the effect of mutations on translation, related to Figure 3A&C.** Selected 5' UTRs (A) or 3' UTRs (B) from the library were ligated with firefly luciferase reporters and co-transfected with *Renilla* luciferase into human cells. The firefly luminescence was normalized to that of *Renilla* (blue bars), reflecting protein output, while RNA expression was measured using quantitative RT-PCR (black bars).

### Figure S5

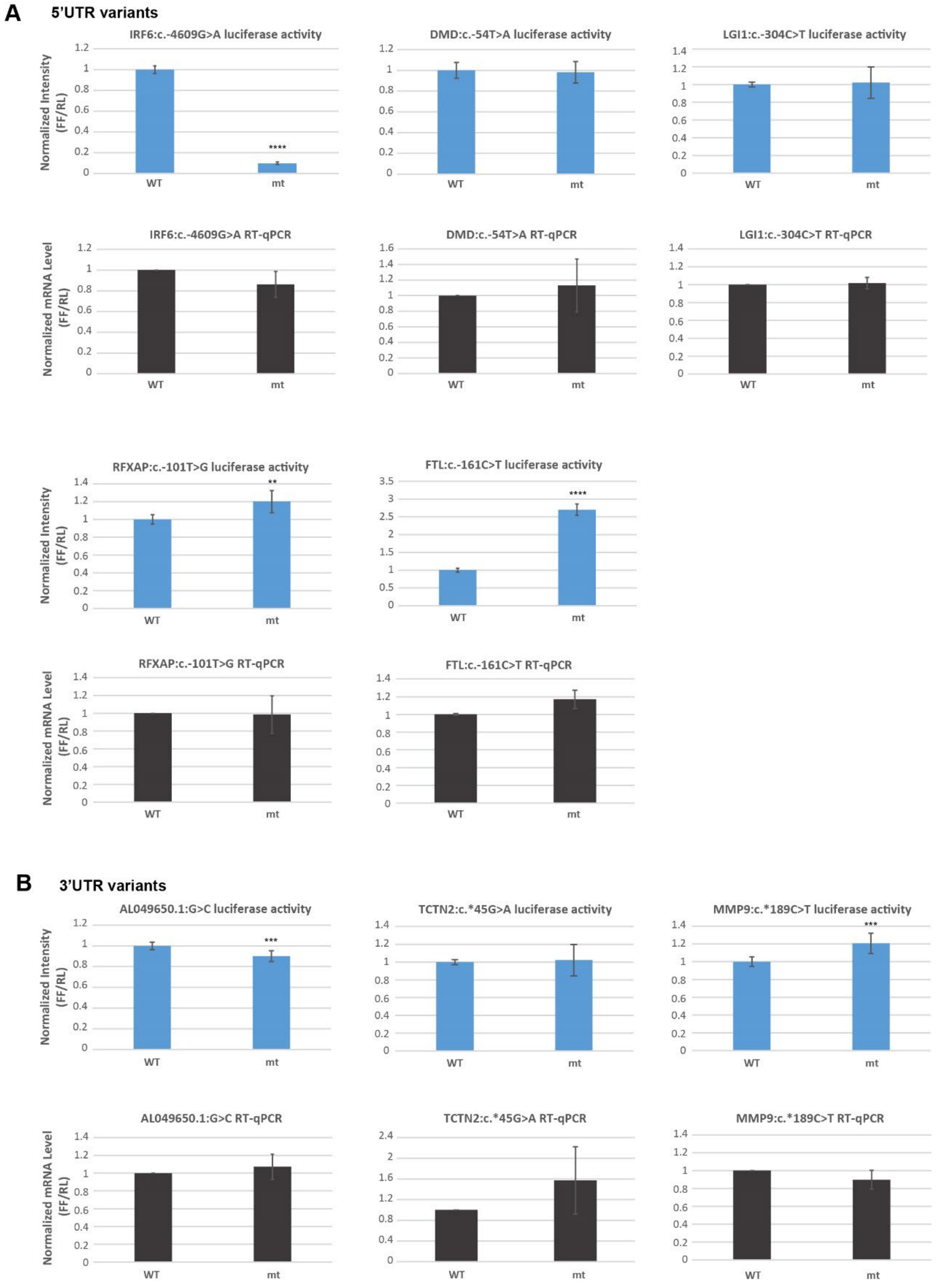


**Figure S5. Validation by luciferase assays of the effect of mutations on translation, related to Figure 3B&C.** Selected full-length 5' UTRs (A) or 3' UTRs (B) were ligated with firefly luciferase reporters and co-transfected with *Renilla* luciferase into human cells. The firefly luminescence was normalized to that of *Renilla* (blue bars), reflecting protein output, while RNA expression was measured using quantitative RT-PCR (black bars).

### Figure S6

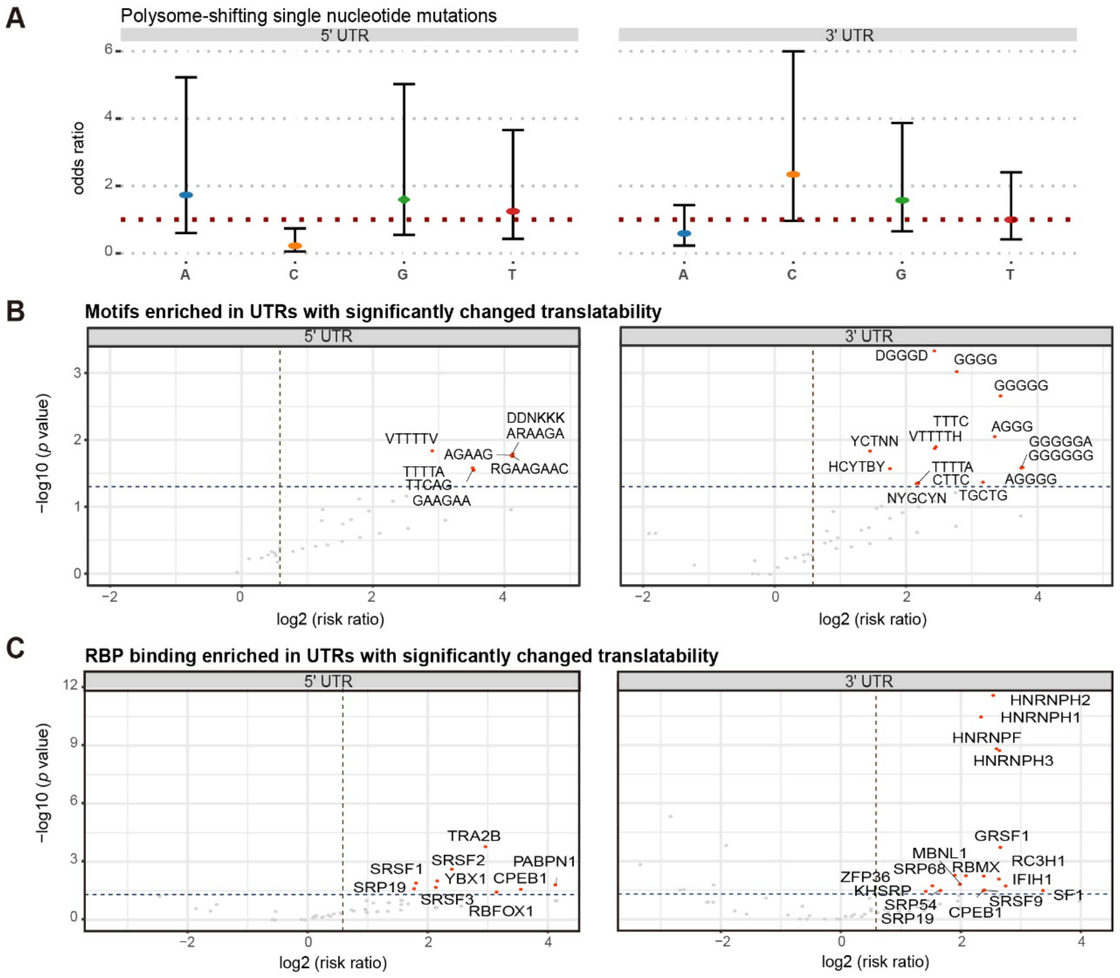


**Figure S6. RBP motif enrichment in polysome-shifting UTRs. (A)** Odds ratios for nucleotide mutations in association with changes in polysome profiles. Error bars represent 95% confidence intervals. Dark red dotted line indicates an odds ratio of 1. Note that alterations of C in 5' UTRs are less likely to prompt a translation defect. **(B)** Motifs enriched in the HC-sig polysome-shifting UTRs. The consensus of the motifs was plotted according to information content. Nucleotide symbols, K: G or T; R: A or G; Y: C or T; V: not T; D: not C; H: not G; B: not A; N: A or C or G or T. **(C)** RBPs enriched among the HC-sig polysome-shifting UTRs.
